# Myosin X interaction with KIF13B, a crucial pathway for Netrin-1-induced axonal development

**DOI:** 10.1101/2020.01.06.896100

**Authors:** Hua-Li Yu, Yun Peng, Yang Zhao, Yong-Sheng Lan, Bo Wang, Lu Zhao, Dong Sun, Jin-Xiu Pan, Zhao-Qi Dong, Lin Mei, Yu-Qiang Ding, Xiao-Juan Zhu, Wen-Cheng Xiong

## Abstract

Myosin X (Myo X) transports cargos to the tip of filopodia for cell adhesion, migration, and neuronal axon guidance. Deleted in Colorectal Cancer (DCC) is one of Myo X cargos essential for Netrin-1-regulated axon pathfinding. Myo X’s function in axon development in vivo and the underlying mechanisms remain poorly understood. Here, we provide evidence for Myo X’s function in Netrin-1-DCC regulated axon development in mouse neocortex. Knocking-out (KO) or knocking-down (KD) Myo X in embryonic cortical neurons impairs axon initiation and contralateral branching/targeting. Similar axon deficits are detected in Netrin-1-KO or DCC-KD cortical neurons. Myo X interacts with KIF13B (a kinesin family motor protein), which is induced by Netrin-1. Netrin-1 promotes anterograde transportation of Myo X into axons in KIF13B dependent manner. KIF13B-KD cortical neurons exhibit similar axon deficits. These results suggest Myo X-KIF13B as a critical pathway for Netrin-1 promoted axon initiation and branching/targeting.

## INTRODUCTION

Neurons are highly polarized cells typically with a single axon and multiple dendrites. Axon development is crutial for the establishement of neuronal connections, especially for the connection between different brain regions. Axon development includes three main steps: (1) axon specification/initiation during neuronal polarization; (2) axon growth and guidance; and (3) axon branching and presynaptic differentiation. During early development of mammalian cortex, the migrating neurons in the intermediate zone (IZ) transit from multipolar to bipolar, with a leading process towards the pia and a trailing process towards the ventricle. Once localized into the cortical plate (CP), the leading processes will develop into highly branched dendrites and the trailing processes will become long axons projecting to target regions for further bifurcation (Barnes and Polleux, 2009; Yogev and Shen, 2017). Eventually, axons connect with target neurons to form synapses that are crucial for neuro-transimission.

Axon development is regulated by intrinsic factors in neurons as well as the micro-envirenmental factors or extracellular guidance cues. Netrin-1 belonges to the netrin family of extracellular guidance cues, which is crucial for axon pathfinding (Colamarino and Tessier-Lavigne, 1995; Braisted et al., 2000). Netrin-1 exerts attractive and repulsive effects through two families of receptors, DCC and UNC-5, respectively (Hedgecock et al., 1990; Ackerman et al., 1997; Leonardo et al., 1997; Culotti and Merz, 1998; Hong et al., 1999; Keleman and Dickson, 2001). Various signaling cascades are involved in Netrin-1-DCC-induced neurite outgrowth and/or growth cone guidance in cultured neurons, which include Rho family GTPases (Li et al., 2002; Shekarabi and Kennedy, 2002), phospholipase C (PLC) (Xie et al., 2006), phosphoinositol 3-kinase (PI 3-kinase) (Ming et al., 1999), extracellular regulated kinase (ERK) (Campbell and Holt, 2003; Forcet et al., 2002; Ming et al., 2002), FAK (Ren et al., 2004; Liu et al., 2004; Li et al., 2004), PITPalpha (Xie et al., 2005), and Myosin X (Myo X) (Zhu et al., 2007). Recent studies using Netrin-1 CKO mice suggest that the ventricular-zone-derived Netrin-1 contributes to commissural axon projection by bounding to commissural axons near the pial surface (Dominici and Moreno-Bravo, et al., 2017). This finding makes the locally produced Netrin-1’s function in commissural axon development more prominent. However, how Netrin-1 regulates cortical neuronal axon development or projection remains to be determined.

Myo X, an unconventional actin based motor protein, is primarily localized at the tips of filopodia or the edges of lamellipodia and membrane ruffles (Berg and Cheney, 2002; Berg et al., 2000; Tokuo and Ikebe, 2004; Zhang et al., 2004). It undergoes forward and rearward movements within filopodia and promotes filopodia formation, elongation, and sensing, possibly by transporting actin binding proteins and cell adhesion receptors to the leading edge of the cells (Berg and Cheney, 2002; Tokuo and Ikebe, 2004; Tokuo et al., 2007; Zhang et al., 2004; Zhu et al., 2007). Myo X has a unique protein structure feature, containing a motor domain at its amino-terminus, three calmodulin-binding IQ motifs, three PH domains, one myosin tail homology (MyTH) domain, and one band 4.1-ezrin-radixin-meosin (FERM) domain (Berg et al., 2000; Yonezawa et al., 2000). Via these domains, Myo X not only binds to F-actin filaments, but also interacts with phosphoinositol lipids, microtubules, and transmembrane receptors (e.g., integrins and DCC family receptors) (Cox et al., 2002a; Plantard et al.; Tokuo and Ikebe, 2004; Weber et al., 2004; Zhang et al., 2004; Zhu et al., 2007). In cultured neurons, Myo X is gradually accumulated to nascent axons, where it regulates axon outgrowth (Yu et al., 2012). In Chicken embryos, expression of motor less Myo X impairs axon growth and commissural axon midline crossing (Zhu et al., 2007). Notice that Myo X is critical for transporting DCC to the dynamic actin-based membrane protrusions (Zhu et al., 2007), and on the other hand, Myo X’s motor activity and distribution are also reciprocally regulated by DCC and neogenin (Liu et al., 2012). Whereas these observations support the view for Netrin-1-DCC-Myo X pathway to be critical for axon development, whether and how they regulate axon development in vivo remain to be further elucidated.

Here, we present evidence for Myo X interaction with KIF13B to be crucial for Netrin-1-induced axon initiation and branching/targeting in developing mouse cortical brain. Myo X interacts with KIF13B (also called GAKIN), a kinesin family member that is essential for delivery of PI(3,4,5)P3 to axons and axon outgrowth (Horiguchi et al., 2006; Yoshimura et al., 2010). Netrin-1 increases Myo X interaction with KIF13B, and thus promoting anterograde transport and axonal targeting of Myo X in neurons. Such Netrin-1’s function requires Myo X interaction with DCC and KIF13B, as well as PI3K activity. Additionally, as Netrin-1 and DCC, both Myo X and KIF13B are required for axon initiation and contralateral branching/targeting in developing cerebral cortex. Taken together, these results suggest that by promoting KIF13B-mediated axonal transport of Myo X, Netrin-1 plays critical roles in inducing axonal initiation and enhancing axonal contralateral branching, revealing unrecognized functions and mechanisms underlying Netrin-1 signaling pathway in axon development.

## RESULTS

### Myo X regulating axonal initiation and contralateral branching/targeting in developing cerebral cortex

To investigate Myo X’s function in axon development in vivo, we used Cre-LoxP recombination technology in combination with in utero electroporation (IUE) to delete Myo X in portions of projection neurons in developing neocortex, and then examined their axon development. Specifically, three processes during axon development, including axon initiation, growth, and branching/targeting, were evaluated as illustrated in Figure 1A. We first evaluated axonal growth and midline crossing in control and MyoX-KO cortical neurons. MyoX-KO cortical neurons were achieved by IUE of Cre-GFP or GFP (as a control) into the neocortex of Myo X^f/f^ embryos (at E15.5) (Wang et al., 2018); and the electroporated brain samples were examined at postnatal day 7 (P7), a critical time window for cortical neuronal axon growth and midline crossing (Figure 1A). To our surprise, axons of Myo X-KO (Cre-GFP^+^) neurons crossed the midline, and their lengths were comparable to those of control axons (Figure 1B), suggesting little role, if there is any, for Myo X to play in axon growth or middling crossing. We second examined axon contralateral branching/targeting in P14 control and Myo X-KO cortical brains. As shown in Figure 1C and 1D, axon branching/targeting to the contralateral cortex was detected in control neurons; but, this process was largely impaired in Myo X-KO neurons. In addition, axon ipsilateral branching was also impaired in the mutant neurons (Figure 1-Figure Supplement 1A and 1B). These results suggest Myo X’s necessity in promoting axon branching/targeting. Third, we accessed Myo X’s function in axon initiation. To this end, Cre-GFP was electroporated into the neocortex of Myo X^f/f^ and wild type (WT) embryos at E14.5, and their axon intensity ratio (defined by Takashi Namba) (Namba et al., 2014) was analyzed at E18.5, a critical time window for axon initiation. As shown in Figure 1E and F, Myo X-KO resulted in a reduction in the axon intensity ratio in the brain, suggesting a role of Myo X in this event. Accordingly, axons in control neurons crossed the midline as early as P3, while axons in MyoX-KD neurons failed to do that (Figure 1-Figure Supplement 1C and 1D). In addition, neuronal migration appeared to be impaired (Figure 1E and G), as reported previously using RNA interference technology (Lai et al., 2015). Given that MyoX KO neurons exhibited normal axonal length and midline crossing at P7, these results suggest a delayed initial axonal outgrowth. Taken together, these results reveal unrecognized roles of Myo X in axon initiation and contralateral branching/targeting in developing cerebral cortex.

**Figure 1.**
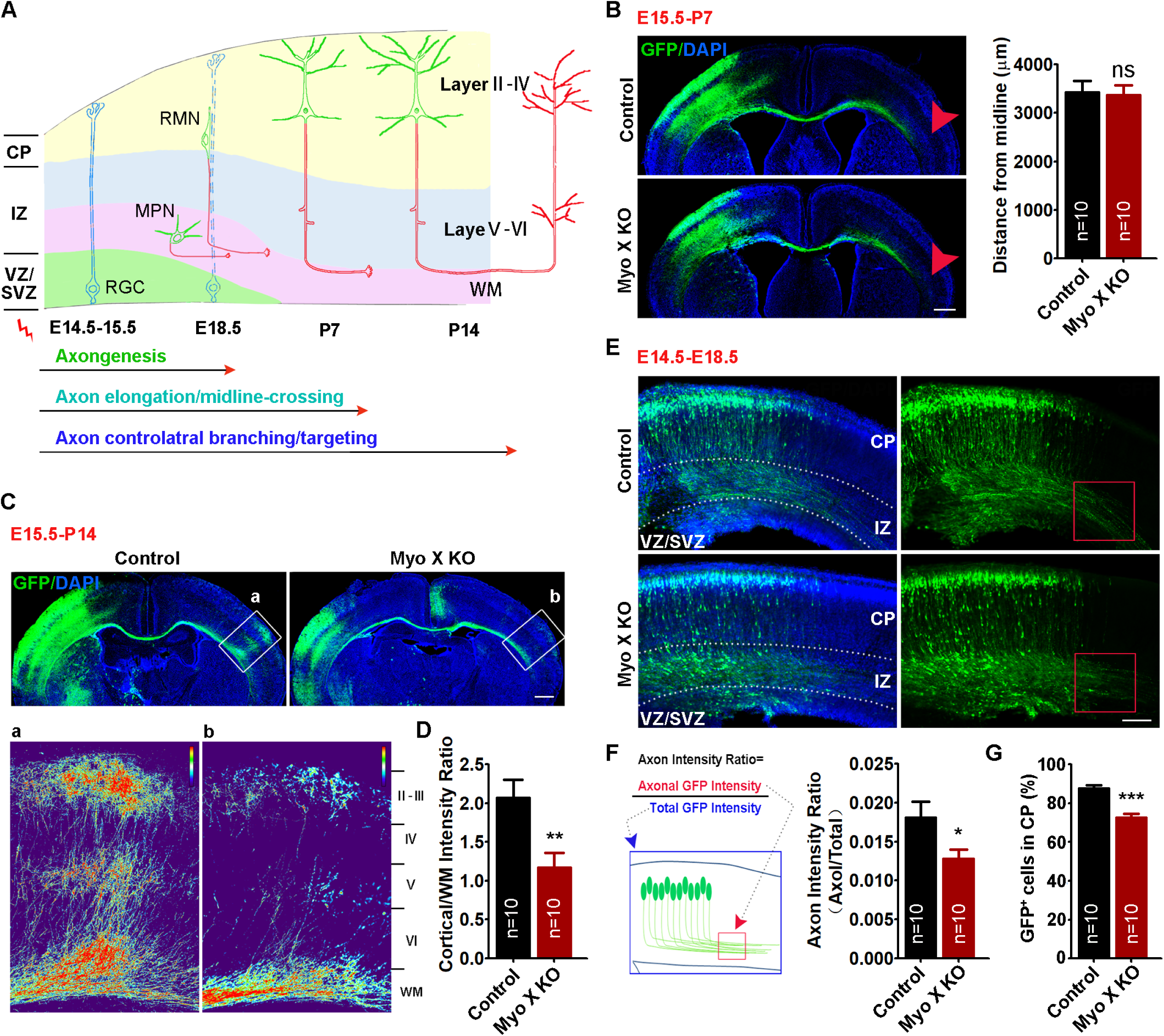
Defective axon initiation and contralateral branching in Myo X-KO cortical neurons. **(A)** Schematic diagram of IUE assay for axon development in neocortex. **(B-D)** P7 (B) and P14 (C-D) cerebral cortex that were electroporated with GFP (Control) or Cre-GFP (Myo X KO) plasmids into Myo X^f/f^ embryos at E15.5. Representative images were shown in B and C. Scale Bar=500μm. Quantifications of axon elongation and axon intensity in the contralateral cortex (as indicated by a and b in C) were presented in B (right panel) and D, respectively. For (B), student’s t test, p = 0.8405; for (D), student’s t test, p = 0.0074. **(E-G)** E18.5 cerebral cortex that were electroporated with Cre-GFP plasmids into Myo X^f/f^ embryos or wild type embryos at E14.5. E, Representative images. Scale Bar=100μm. F, Quantification of axon initiation by using the axon intensity ratio, which is defined as axonal GFP intensity (marked by a red square) over the total GFP intensity (marked as a blue square), as illustrated in the schematic diagram. Student’s t test, p = 0.0379. G, Quantification of GFP^+^ cells in CP. Student’s t test, p<0.0001. Data are presented as the means ± SEM. The numbers of brain sections scored are from 3 different brains for each group and indicated on the graphs. ns, no significant difference; *, P<0.05; **, P<0.01; ***, P<0.001.

### Netrin-1 promoting Myo X-regulated axonal initiation and branching

Given the specific orientation of the trailing processes or nascent axons, we speculate that extracellular signals from ventricular zone and sub-ventricular zone (VZ/SVZ) modulate intrinsic signaling (e.g., Myo X) for axon initiation and development, and Netrin-1, an upstream Myo X regulator (Zhu et al., 2007), may be involved in axon initiation and branching. To test this speculation, we examined Netrin-1’s function in axon development by IUE of Cre-GFP into Netrin-1 floxed (NTN1^f/f^) embryos at E14.5 (Figure 2-Figure Supplement 1A). The electroporated brain samples were examined at E18.5 to evaluate the neuronal axon intensity ratio. Indeed, a reduction in the axon intensity ratio of cortical neurons was detected in Netrin-1 KO embryos (Figure 2A and 2B), revealing a similar role of Netrin-1 as that of Myo X in axon initiation. Neuronal migration was not affected by Netrin-1 KO (Figure 2A and 2C). We next asked if Netrin-1-regulated axon initiation depends on Myo X. To this end, plasmids encoding Myc-Netrin-1 and Myo X miRNA were co-electroporated into the E14.5 embryos (Figure 2D). Netrin-1 ectopic expression restored the axon intensity ratio in Myo X-KD neurons (Figure 2D and 2E), supporting the view for Netrin-1-Myo X pathway in promoting axon initiation.

**Figure 2.**
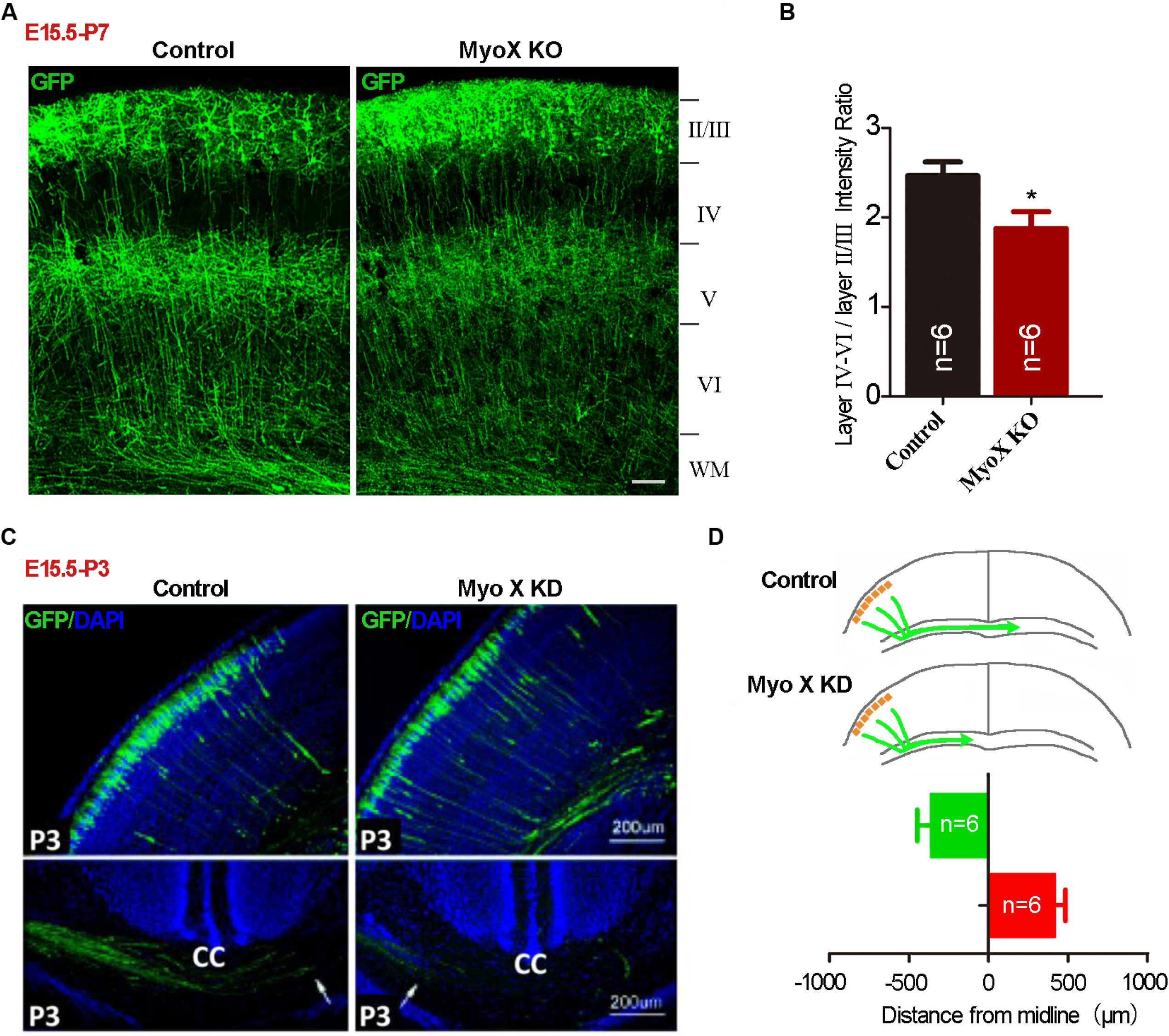
Netrin-1 rescue of axon initiation defect caused by Myo X deficiency. **(A-C)** E18.5 cerebral cortex that were electroporated with GFP (Control) or Cre-GFP (Netrin-1 KO) plasmids into NTN1^f/f^ embryos at E14.5. A, Representative images. Scale Bar=100μm. B, Quantification of axon intensity ratio. Student’s t test, p = 0.0425. C, Quantification of GFP^+^ cells in CP. Student’s t test, p =0.3970. **(D)** Representative images of E18.5 cerebral cortex electroporated with Control miRNA (Control), Myo X miRNA (Myo X KD) or Myo X miRNA together with Myc-Netrin-1 plasmids (Myo X KD + Netrin-1) at E14.5. Scale Bar=100μm. **(E)** Quantification of axon intensity ratio. One-way ANOVA, p =0.0303 for MyoX KD group, p=0.7385 for Myo X KD+Netrin-1 group. **(F)** Representative images of P14 cerebral cortex electroporated with Control miRNA (Control), Netrin-1 shRNA (Netrin-1 KD), DCC miRNA (DCC KD) at E15.5. Scale Bar=500μm. **(G)** Quantification of axon contralateral branching. Student’s t test, p =0.398. Data are presented as the means ± SEM. The numbers of brain sections scored are from 3 different brains for each group and indicated on the graphs. ns, no significant difference; *, P<0.05; **, P<0.01.

We then asked whether Netrin-1-DCC pathway play a role in axon branching/targeting as Myo X does. Netrin-1 or DCC expression in E15.5 cortical neurons were suppressed by IUE of their shRNAs, respectively. Their axons at age of P14 were examined. Netrin-1-KD in E15.5 cortical neurons had little effect on the axonal contralateral branching/targeting (Figure 2F and 2G) or ipsilateral branching (Figure 2-Figure Supplement 1B and 1C). However, DCC KD impaired axonal contralateral branching/targeting (Figure 2F) as well as ipsilateral branching (Figure 2-Figure Supplement 1B and 1C). These results, in line with our hypothesis, suggest that DCC and Myo X in neurons are necessary to promote axon branching, but, Netrin-1, an extracellular cue, regulates axon development in cell non-autonomous fashion.

### Netrin-1 increasing axonal distribution and transport of Myo X in cultured neurons

To understand how Netrin-1 regulates Myo X’s function in axon development, we examined Netrin-1’s effect on exogenous Myo X (GFP-Myo X) distribution in cultured neurons. To our surprise, GFP-Myo X was largely distributed in the soma and the tips of dendritic like filopodia, but nearly undetectable in Tau-1 positive axonal compartments in the absence of Netrin-1(Figure 3A). Upon Netrin-1 stimulation, an obvious increase of GFP-Myo X in Tau-1 positive axons with a slight decrease of GFP-Myo X in MAP2 positive dendritic neurites were detected (Figure 3A, 3B, 3C and 3D), suggesting a role of Netrin-1 in regulating GFP-Myo X distribution in axons and dendrites.

**Figure 3.**
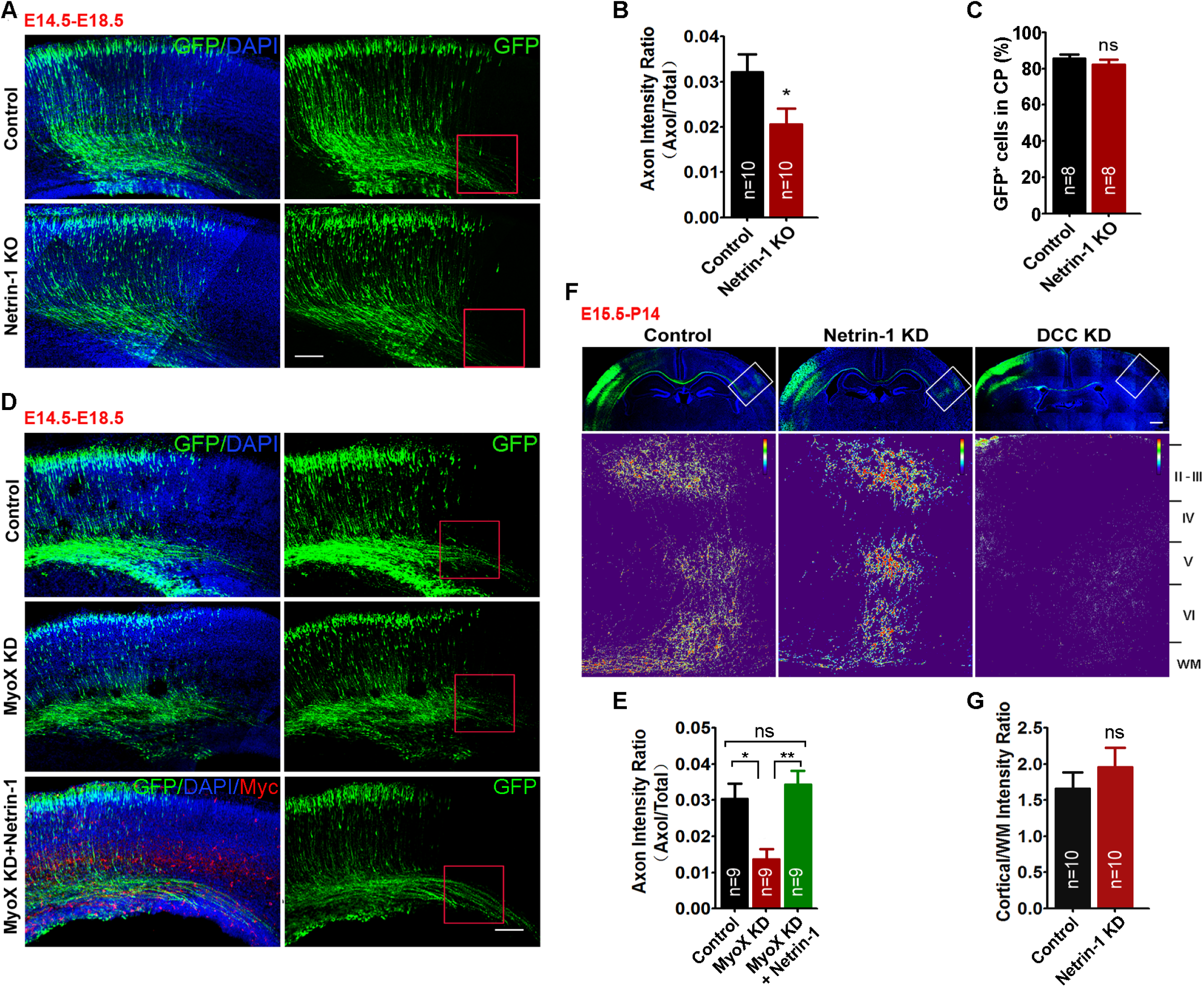
Increase of axonal distribution and anterograde transport of GFP-Myo X by Netrin-1. **(A-D)** Immunostaining analysis using indicated antibodies at DIV 3 neurons that were transfected with GFP-Myo X at DIV 1 and treated with vehicle or Netrin-1 for 1 hr. A, Images marked in rectangular were amplified and showed in the right panels. White arrow heads indicate the axonal distribution of GFP-Myo X. Scale Bar=10μm. B, Quantification of GFP-Myo X intensity in axons. The axonal GFP-Myo X level was normalized to somatic GFP-Myo X. Student’s t test, p = 0.0012. C, Yellow arrows indicate GFP-Myo X distribution in MAP2 positive dendrites. Scale Bar=10μm. D, Quantification of GFP-Myo X intensity in dendrites. The dendritic GFP-Myo X level was normalized to somatic GFP-Myo X. Student’s t test, p = 0.0471. **(E-G)** Analysis of axonal GFP-Myo X mobility with fluorescence recovery after photobleaching (FRAP) assay. E, Images of FRAP analysis in GFP-Myo X expressing neurons in the presence of vehicle or Netrin-1 and the quantification (normalized GFP-Myo X intensity of the photobleached axon compartment). Scale Bar=20μm. F, Quantification of half-time of maximum recovery (t1/2). Student’s t test, p = 0.0446, n=6 neurons from 3 different experiments. G, Percentage of GFP-Myo X recovery. Student’s t test, p<0.0001, n=6 neurons from 3 different experiments. **(H-J)** Time-lapse imaging analysis of GFP-Myo X expressing neurons in the presence of vehicle or Netrin-1. H, the mCherry was co-transfected to visualize neuronal processes. Images marked in rectangular were amplified and showed in the middle panels. Mobile trajectory of indicated GFP-Myo X puncta was presented in the bottom panels. Scale Bar=20μm. I, Quantification of mean velocity of GFP-Myo X puncta. Student’s t test, p <0.0001. J, Quantification of stationary GFP-Myo X. Student’s t test, p = 0.021. Data are presented as the means ± SEM. The numbers of neurons scored are from 3 different experiments for each group and indicated on the graphs. *, P<0.05; **, P<0.01; ***, P<0.001.

GFP-Myo X in axons appeared to be diffusible (Figure 3A), exhibiting the characteristics of slow anterograde transport (Brown, 2003; Maday et al., 2014). We thus examined the dynamics of GFP-Myo X in axons by fluorescence recovery assay after photo bleaching (FRAP) (Figure 3E). The fluorescence recovery of GFP-Myo X in control axons was much slower and incomplete than that of Netrin-1 treated axons (Figure 3E, 3F and 3G), supporting the view for Netrin-1 to enhance GFP-Myo X movement. In contrast from axonal GFP-Myo X, GFP-Myo X in dendrite-like filopodia exhibited puncta pattern (Figure 3H). Time-lapse imaging and analyzing the motility of GFP-Myo X in these filopodia showed both extension and retraction movements of GFP-Myo X puncta in control neurons (Figure 3H, bottom panels). Upon Netrin-1 stimulation, the travelling path and the average velocity of GFP-Myo X puncta were all reduced (Figure 3H, bottom panels, and 3I), with an increase in the percentage of stationary puncta (Figure 3J). Together, these results suggest that while Netrin-1 increases anterograde movement of GFP-Myo X in axons, it decreases GFP-Myo X’s motility in dendrite-like filopodia.

### Requirement of DCC and PI3K activation for Netrin-1-increased axonal distribution of Myo X

To further understand how Netrin-1 regulates Myo X’s axonal distribution, we first mapped domains in GFP-Myo X that are necessary for this event. Myo X is a multi-domain containing unconventional myosin, containing 3 PH domains, a myosin tail homology 4 (MyTh4) domain, and a band 4.1–ezrin–radixin–moesin (FERM) domain in the C-terminus, in addition to the motor domain in the N-terminus (Berg et al., 2000; Kerber and Cheney, 2011). GFP-Myo X deletion mutants were generated as illustrated in Figure 3-Figure Supplement 1A (left panel). In the absence of Netrin-1, Myo X mutants containing the motor domain showed a similar distribution pattern as that of full length Myo X, with predominant localizations in the soma and dendritic filopodia (Figure 3-Figure Supplement 1A and 1B). However, in the presence of Netrin-1, Myo X deletions in the second PH domain (Myo X^ΔPH2^) or in the FERM domain (Myo X^ΔFERM^) abolished Netrin-1-induced axonal distribution (Figure 3-Figure Supplement 1A and 1B). These results suggest a requirement of both PH and FERM domains in Myo X for Netrin-1-induced Myo X’s axonal distribution.

As the FERM domain in Myo X binds to DCC (Zhu et al., 2007), we thus speculate that DCC-Myo X interaction may be critical for this event. Indeed, this view is supported by the observations that DCC-Myo X interaction is up-regulated by Netrin-1 (Figure 3-Figure Supplement 2A and 2B), and neurons suppressing DCC expression by its miRNA failed to response to Netrin-1 in targeting Myo X to axons (Figure 3-Figure Supplement 2C, 2D and 2E).

Myo X’s PH domains are known to bind to multi-phosphoinositols, including PI(4,5)P2, PI(3,4)P2, and PI(3,4,5)P3 (Cox et al., 2002; Umeki et al., 2011). The second PH domain is crucial for binding to PI(3,4,5)P3/PI(3,4)P2, products of PI3K activation (Umeki et al., 2011). As this PH domain in Myo X is necessary for Netrin-1-induced axonal distribution of Myo X, we wondered if PI(3,4,5)P3/PI(3,4)P2 are involved in Myo X’s axonal distribution. To this end, neurons expressing GFP-Myo X were treated with Netrin-1 in the presence or absence of wortmannin (10 nM), an inhibitor of PI3K that blocks production of PI(3,4,5)P3 and PI(3,4)P2. Such an inhibition abolished Netrin-1-induced Myo X’s axonal distribution (Figure 3-Figure Supplement 2F, 2G and 2H), supporting a role for PI3K activation and its products, PI(3,4,5)P3/PI(3,4)P2, in this event.

### Myo X interaction with KIF13B, a kinesin family motor protein

Given that anterograde transport is powered by kinesin family motor, and Myo X binds to microtubules (Weber et al., 2004), we asked if Myo X acted as a cargo of kinesin motor protein to be transported along microtubules in axons. Immunoprecipitation assay was used to screen for GFP-Myo X-binding kinesins, including KIF1B, KIF3A, KIF3C, KIF5 and KIF13B/GAKIN, which are well recognized kinesin motor proteins in axonal anterograde transportation (Hirokawa, 2009). Interestingly, KIF13B, which is essential for anterograde transport of PI(3,4,5)P3 for axonal outgrowth and formation (Yoshimura et al., 2010), was detected in the GFP-Myo X immunoprecipitates in primary cultured neuronal lysates (Data not shown).

We then mapped the domains in KIF13B for its interaction with Myo X by coimmunoprecipitation assay. KIF13B contains a motor domain at the NH2 terminus, a forkhead-associated (FHA) domain, a MAGUK binding stalk (MBS) domain, two domains of unknown function (DUF) and a CAP-Gly motif at the COOH terminus (Figure 4A). As shown in Figure 4A, C terminal regions of KIF13B (Myc tagged) (including KIF13B^558-1826^, KIF13B^990-1826^ and KIF13B^1532-1826^), but not the N-terminus (KIF13B^1-557^), were detected in Myo X immunoprecipitates. Further analysis of their interaction identified that the C-terminal domain, KIF13B^1532-1826^, is involved in the interaction with Myo X.

**Figure 4.**
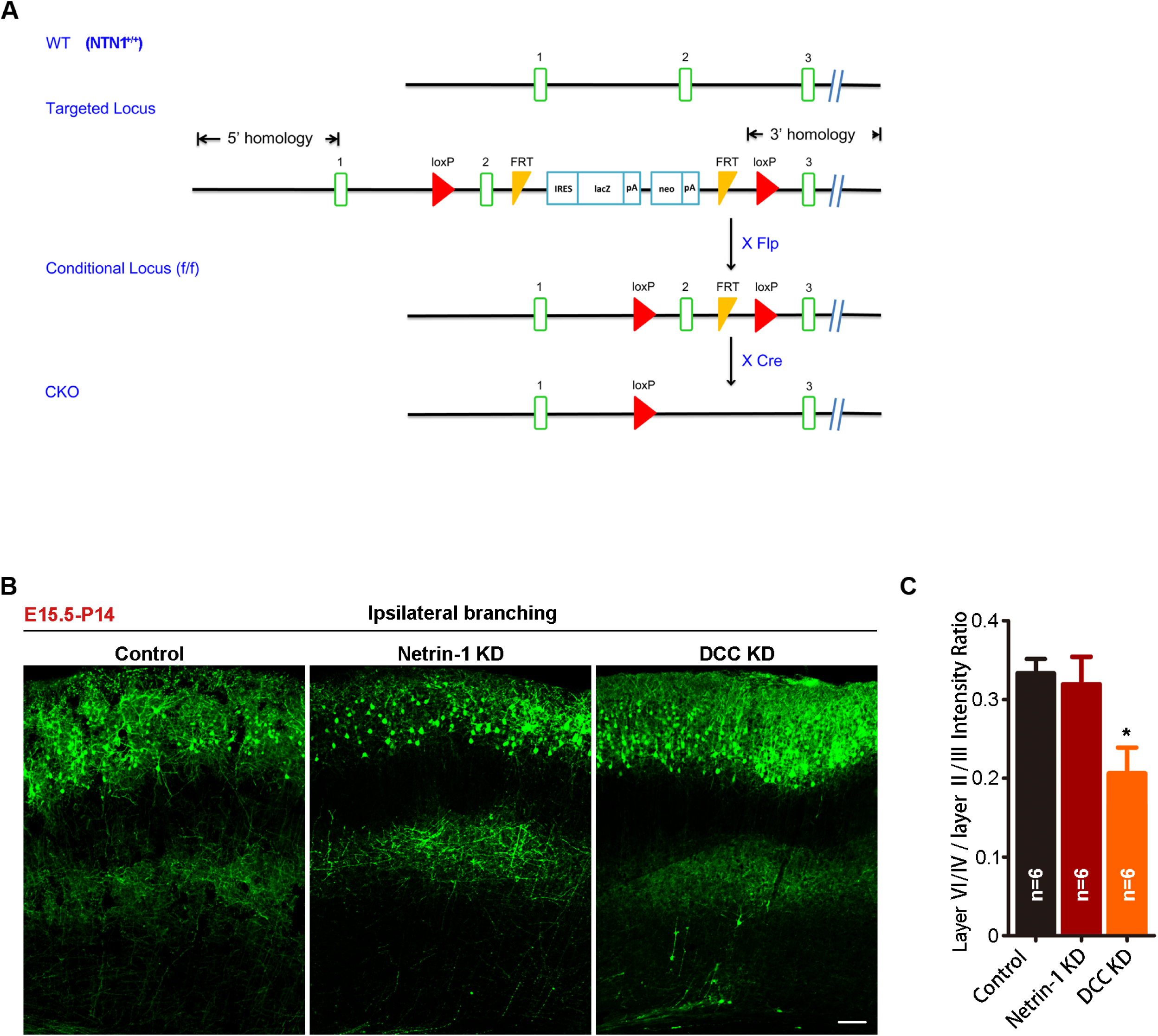
Interaction of Myo X with KIF13B. **(A)** Co-immunoprecipitation of GFP-hMyo X and Myc-KIF13B^1532-1826^. GFP-hMyo X was co-expressed with Myc-KIF13B and its deletion mutants in HEK293T cells and immunoprecipitated by anti-Myc antibody. **(B)** Immunoblotting of the pulled down fraction by the GST-KIF13B C-terminus fusion protein and GST alone. The amounts of GST fusion proteins and GST were revealed by coomassie blue staining (lower panel). **(C)** Immunoblotting of the pulled down fraction by the truncated C-terminal domains fused with GST and GST alone. The amounts of different GST fusion proteins and GST alone were revealed by coomassie blue staining (lower panel). **(D)** Co-localization of GFP-Myo X and Myc-KIF13B in filopodia tips of NLT cells (upper panels) and in axon and dendrite-like filaments in cultured cortical neurons (lower panels). Images marked in rectangular was amplified and included as insert. Scale Bar=20μm. **(E)** Co-immunoprecipitation of exogenous Myc-KIF13B and GFP-Myo X. HEK293T cell lysates were immunoprecipitated with anti-Myc antibody and with IgG as control. **(F)** Immunoprecipitation of endogenous Myo X and KIF13B with or without Netrin-1 stimulation. Neurons treated with vehicle or Netrin-1 were lysed and incubated with anti-Myo X antibody (upper panel). Quantifications of KIF13B binding with MyoX were presented in lower panel. student’s t test, p = 0.0089. Data are presented as the means ± SEM. For statistical analysis, three independent experiments were performed and indicated on the graphs. **, P<0.01.

The Myo X-KIF13B interaction was further verified by a glutathione S-transferase (GST) pulldown assay. The recombinant GST-KIF13B^1532-1826^ fusion protein was produced (Figure 4B), which was used to pull down lysates expressing various GFP-Myo X mutants (including Myo X^Head^, hMyo X, hMyo X^ΔPH2^, hMyo X^ΔPH3^, hMyo X^KK1225/6AA^, Myo X^Myth4-Ferm^ and Myo X^Ferm^). hMyoX is an abbreviation of headless MyoX, which contains amino acids from 860 to 2062. hMyoX^KK1225/6AA^ means that the 1225/1226 Lysine was further mutated to Alanine. These two lysines are located in the second PH domain and required for MyoX binding with PI(3,4,5)P3. Note that only hMyo X and hMyo X^KK1225/6AA^ were pulled down by GST-KIF13B^1532-1826^, suggesting the requirement of the second and third PH domains of Myo X for its binding to the C-terminal domain in KIF13B (Figure 4B). By this assay, the effective Myo X binding region in KIF13B was further mapped to the last 74 amino acids in its C-terminus (Figure 4C). It is noteworthy that while the site KK1225/6 in Myo X is critical for binding to PI(3,4,5)P3 (Plantard et al., 2010), it was not required for Myo X interaction with KIF13B.

Finally, we examined Myo X-KIF13B interaction by co-immunostaining analysis. As shown in Figure 4D, GFP-Myo X was co-localized with Myc-KIF13B in filopodia tips in NLT cells and in axon and dendrite-like filaments in cultured cortical neurons. In addition, their interaction was reconfirmed by co-immunoprecipitation analysis of exogenously expressed GFP-Myo X and Myc-KIF13B (Figure 4E), as well as endogenous KIF13B with Myo X in primary neuronal lysates (Figure 4F). Interestingly, Netrin-1 stimulation increased Myo X-KIF13B interaction in neurons (Figure 4F). Taken together, these results suggest that Myo X interacts with KIF13B, implicating KIF13B in Netrin-1-induced Myo X distribution in axons.

### Myo X as a cargo of KIF13B for its axonal distribution

To investigate KIF13B’s function in Netrin-1 induced Myo X anterograde transportation, we first examined whether GFP-Myo X’s distribution in Tau-1 positive axons was affected by KIF13B expression. Indeed, expression of KIF13B was sufficient to increase GFP-Myo X’s localization in axons and decrease GFP-Myo X’s localization in dendrites in the absence of Netrin-1(Figure 5A, 5B and 5C). In line with this view was the observation by the FRAP assay that the recovery of GFP-Myo X after photo-bleaching in axons was speed up by KIF13B expression (Figure 5D, 5E and 5F). Furthermore, KIF13B’s effect on GFP-Myo X distribution was examined by time lapse imaging analysis. As shown in Figure 5G, GFP-Myo X puncta exhibited high motility in both dendrite-like neurites and growth cones. Such actin-based motility of GFP-Myo X was decreased in neurons co-expressing KIF13B (Figure 5G, 5H and 5I).

**Figure 5.**
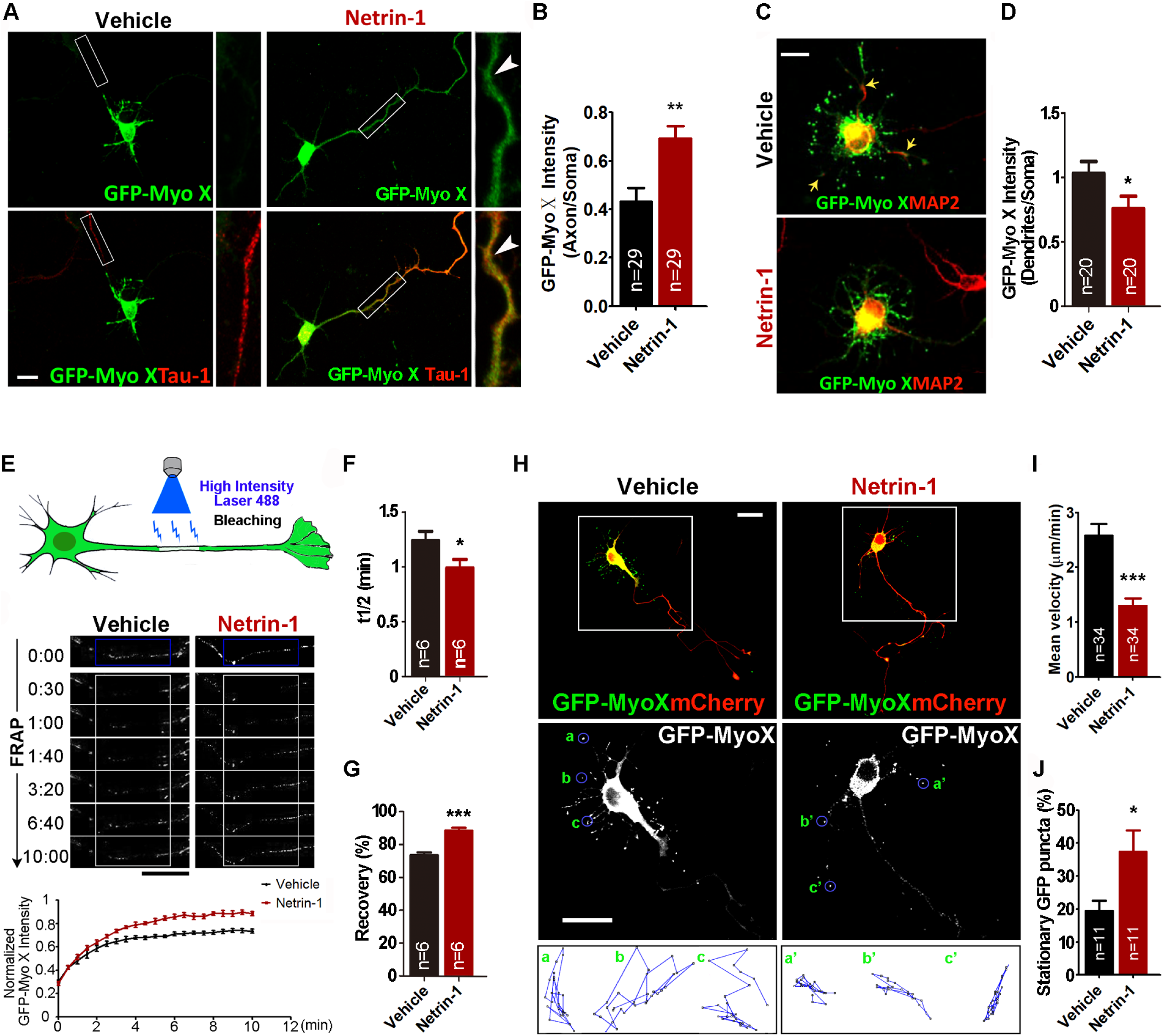
Involvement of KIF13B in Myo X anterograde transportation. **(A-C)** Immunostaining analysis using indicated antibodies at DIV 3 neurons that were transfected with GFP-Myo X or GFP-Myo X together with Myc-KIF13B at DIV 1. A, Images marked in rectangular were amplified and showed in the right panels. Scale Bar=20μm. B, Quantification of GFP-Myo X intensity in axons. The axonal GFP-Myo X level was normalized to somatic GFP-Myo X. Student’s t test, p <0.0001. C, Quantification of GFP-Myo X intensity in dendrites. The dendritic GFP-Myo X level was normalized to somatic GFP-Myo X. Student’s t test, p=0.0013. **(D-F)** Analysis of KIF13B effect on axonal GFP-Myo X mobility with FRAP assay. D, Images of FRAP analysis in DIV 3 cortical neurons that were transfected with GFP-Myo X or GFP-Myo X together with Myc-KIF13B at DIV 1 and the quantification (normalized GFP-Myo X intensity of the photobleached axon compartment). Scale Bar=20μm. E, Quantification of half-time of maximum recovery. Student’s t test, p=0.0213, n=6 neurons from 3 different experiments. F, Percentage of GFP-Myo X recovery. Student’s t test, p<0.0001, n=6 neurons from 3 different experiments. **(G-I)** Time-lapse imaging analysis of GFP-Myo X together with vector or GFP-Myo X together with Myc-KIF13B transfected neurons. G, The marked rectangular in G were further analyzed by kymographs (see lower panels), which show the mobility of GFP-Myo X positive vesicles during 5-min recordings. Vertical lines represent stationary Myo X-vesicles; oblique lines or curves to the right represent anterograde movements and lines to the left indicate retrograde transport. Scale Bar=20μm. H, Quantification of mean velocity of GFP-Myo X puncta. Student’s t test, p = 0.0026. I, Quantification of stationary GFP-Myo X. Student’s t test, p = 0.0001. Data are presented as the means ± SEM. The numbers of cells scored are from 3 different brains for each group and indicated on the graphs. *, P<0.05; **, P<0.01; ***, P<0.001.

Second, we determined whether KIF13B was necessary for Netrin-1-induced Myo X axonal distribution. Plasmid encoding KIF13B shRNA was generated, which selectively suppressed KIF13B expression (Figure 5-Figure Supplement 1A and 1B). GFP-Myo X with KIF13B shRNA or control shRNA were co-transfected into neurons treated with or without Netrin-1. In control neurons (GFP-Myo X with control shRNA), GFP-Myo X’s distribution in axons was increased and in dendrites was decreased by Netrin-1 stimulation (Figure 5-Figure Supplement 1C, 1D and 1E). However, such an increase of Myo X’s axonal distribution was abolished in neurons co-transfected with KIF13B shRNA (Figure 5-Figure Supplement 1C and 1D). Moreover, KIF13B-KD in embryonic mouse cortical neurons significantly suppressed GFP-Myo X distribution in axon-like neurites (Figure 5-Figure Supplement 1F and 1G). Taken together, these results suggest that KIF13B is not only sufficient, but also necessary for Netrin-1-induced Myo X distribution in axons. In line with this view, KIF13B’s distribution in axons was increased by Netrin-1 (Figure 5-Figure Supplement 1H, 1I and 1J).

### KIF13B, as Myo X, promoting axonal initiation and contralateral branching/targeting in developing cerebral cortex

The important role of KIF13B in Netrin-1 induced Myo X axonal distribution led us to speculate a similar role that KIF13B plays as that of Myo X in Netrin-1-induced axonal initiation and targeting in vivo. To test this speculation, KIF13B shRNA and Myo X miRNA were IUEed into the progenitor cells of cortical neurons in E14.5 mouse embryos, and their brain sections at E18.5 were examined (Figure 6A). As shown in Figure 6A and 6B, KIF13B-KD resulted in a decrease in axon intensity ratio in the cortical brains, a similar impairment in axon initiation as that of Myo X-KD neurons. Besides, the percentage of Myo X or KIF13B deficient neurons into CP was decreased, as compared with that of control neurons (Figure 6A and 6C). Furthermore, we analyzed the polarization of neurons in IZ and defined the longest neurites as axons by two criteria: the length of longest neurite is >50 μm; and 2 times more than that of the second longest one. Based on these criteria, the percentages of polarized neurons in both Myo X-KD and KIF13B-KD groups were decreased (Figure 6D and 6E) and their axons were also shorter (Figure 6D and 6F)). These results suggest that KIF13B plays a similar role as that of Myo X in axon initiation.

**Figure 6.**
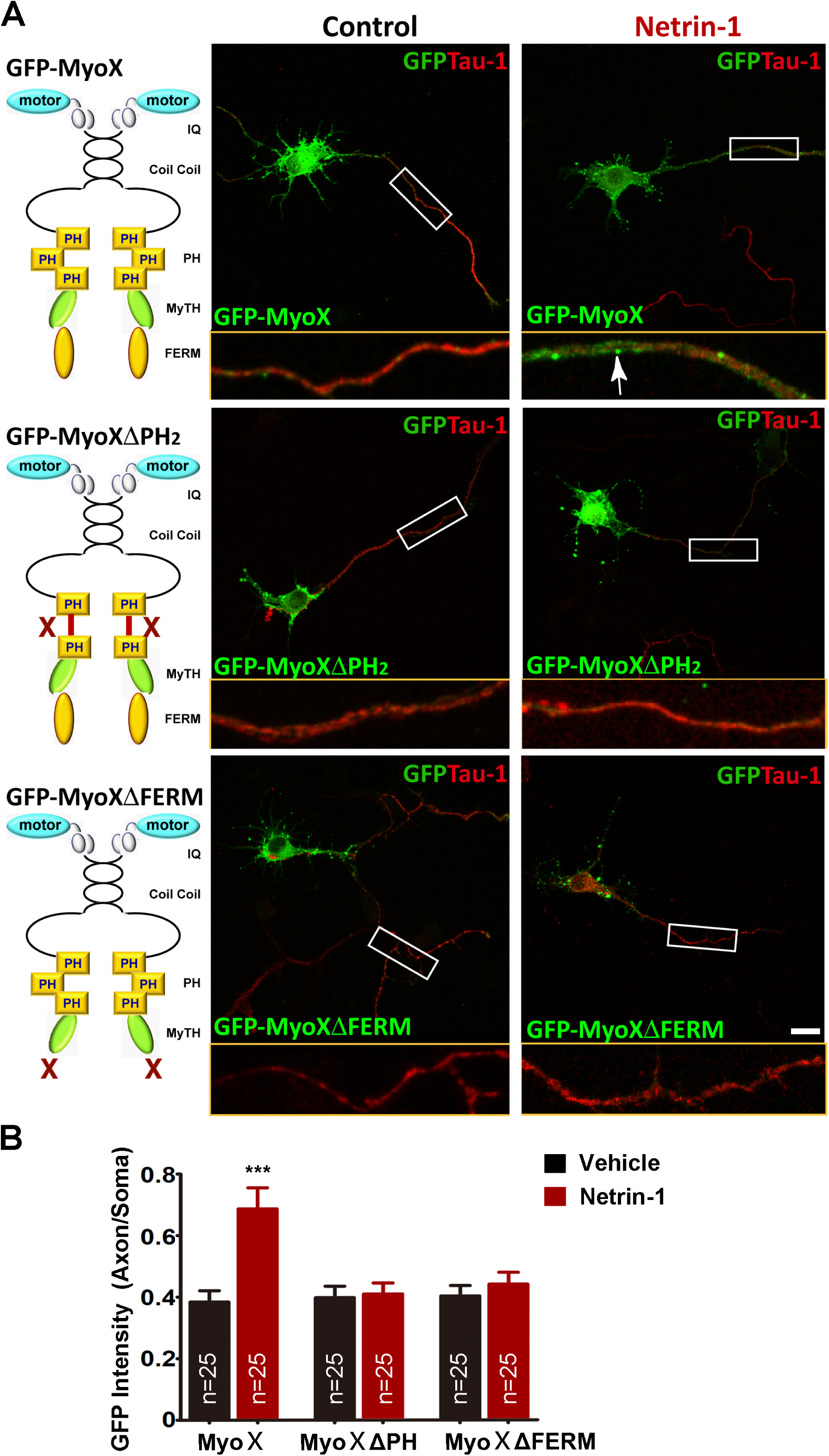
Similar role of KIF13B in axon initiation and branching as Myo X. **(A)** Representative images of E18.5 cerebral cortex electroporated with control shRNA (Control), KIF13B shRNA (KIF13B KD) and Myo X miRNA (Myo X KD) plasmids at E14.5. Scale Bar=100μm. **(B)** Quantification of axon initiation by using the axon intensity ratio. Student’s t test, for Myo X KD group, p=0.0150, for KIF13B KD group, p=0.0102. **(C)** Quantification of GFP^+^ cells in CP. Student’s t test, for Myo X KD group, p=0.0002, for KIF13B KD group, p<0.0001. **(D)** Neurons in the IZ of E18.5 cerebral cortex in each group. Mouse cortices electroporated with indicated plasmids were presented in the upper panels. Tracing of representative GFP+ neurons in each group were presented in the lower panels. Scale Bar=100μm. **(E)** Quantification of polarized neurons in IZ. Student’s t test, for Myo X KD group, p<0.0001, for KIF13B KD group, p<0.0001. **(F)** Quantification of the length of longest neurites. Student’s t test, for Myo X KD group, p=0.0003, for KIF13B KD group, p=0.0003. **(G-H)** Representative images of P7 cerebral cortex electroporated with above indicated plasmids at E15.5 as well as quantification of axon elongation. Scale Bar=500μm. Student’s t test, for Myo X KD group, p=0.9059, for KIF13B KD group, p=0.6466. **(I-J)** Representative images of P14 cerebral cortex electroporated with above indicated plasmids at E15.5 as well as quantification of axon contralateral branching. Scale Bar=100μm. Student’s t test, for Myo X KD group, p=0.0013, for KIF13B KD group, p=0.0067. Data are presented as the means ± SEM. The numbers of brain sections or cells scored are from 3 different brains for each group and indicated on the graphs. ns, no significant difference; *, P<0.05; **, P<0.01; ***, P<0.001.

We next determined if KIF13B regulates axon projection and branching, as Myo X does. To this end, KIF13B was suppressed in E15.5 embryos by IUE of its shRNA (GFP). At neonatal age (e.g., P7 and P14), the IUEed neurons were mostly migrated into cortical L2/3 pyramidal neurons whose axons project to the contralateral side via corpus callosum (CC) (Alcamo et al., 2008). As that of Myo X-KD axons, axons of KIF13B-KD neurons crossed the midline at P7, without an obvious reduction in their axonal length (Figure 6G and 6H). However, at P14, the axonal contralateral branches were severely diminished in KIF13B-KD neurons, compared with that of controls (Figure 6I and 6J). Taken together, these results suggest that KIF13B is necessary to promote axon initiation and branching/targeting in developing cortical neurons, providing additional support for KIF13B as an important mediator for Netrin-1-induced and Myo X-regulated axon initiation and targeting.

## DISCUSSION

In this study, we present evidence that Netrin-1 increases axonal targeting of Myo X in neurons. This event is essential for axon initiation and contralateral branching, but not midline-crossing. Further mechanical studies suggest that Netrin-1 increases Myo X interaction with KIF13B, thus promoting axonal transport of Myo X, axonal initiation and branching/targeting. These results reveal a new mechanism underlying Netrin-1-regulated axon pathfinding.

As an unconventional Myosin family protein, Myo X is widely expressed and implicated in multiple cellular functions in different cell types, including Netrin-1-induced neurite outgrowth and growth cone guidance) (Zhu et al., 2007), BMP6-dependent filopodial migration and activation of BMP receptors (Pi et al., 2007), and migration of *Xenopus* cranial neural crest cells (Hwang et al., 2009; Nie et al., 2009). While it is evident that Myo X modulates growth cone actin dynamics and promotes axon specification in cultured neurons (Yu et al., 2012), the in vivo evidence for this view is limited. Here, we found that Myo X is critical for axon initiation and terminal branching/targeting in developing neocortex. Myo X KD (by RNA interference) or KO (by Cre-Loxp Combination) decreased axon intensity ratio, suggesting a deficit in axon initiation (Figure 1E and 6A). Morphology analysis of individual neurons suggested an impairment of axon genesis (Figure 6D, 6E and 6F). In addition to axon initiation, Myo X KD or KO also impaired axon terminal branching/targeting (Figure 1C and 6I). This Myo X’s function is in line with the observations that Myo X plays an important role in regulating axon filaments and actin filaments in the leading margin of growth cones which are responsible for perceive extrinsic guidance factors or adhesive signals and producing traction for axon terminal elongation towards its target (Yu et al., 2012; Dent and Gertler, 2003). Interesting, Netrin-1 KD in cortical neurons had little effect on the axonal contralateral branching/targeting, while DCC KD impaired axonal contralateral branching/targeting (Figure 2F and 2G). These results together suggest that DCC and Myo X in neurons play a cell autonomous role in promoting axon branching, while Netrin-1, as an extracellular cue, regulates axon development in a non-autonomous way. Although Netrin-1-DCC-Myo X was involved in axon branching, it is not the only pathway underlying axon branching. It is of interest to note that Netrin-1 promotes exocytosis and plasma membrane expansion for axon branching via TRIM9 release of SNAP25 and SNARE-mediated vesicle fusion (Winkle et al., 2014). It will be of interest to investigate if Myo X is involved in Netrin-1 stimulated exocytosis for axon branching. Notice that Myo X-KO or KD has little effect on axon midline crossing, so does in Netrin-1-KO or DCC-KD axons (Figure 2F and 2G). These results suggest a neuronal DCC-Myo X independent mechanism for axon midline crossing.

Axons contain abundant microtubules, although their growth cones have enriched actin filaments (Dent and Gertler, 2003). How is Myo X, an actin-filament based motor protein, transported to the growth cones of axons? Although Myo X interacts with microtubules with its MyTH4 domain (Weber et al., 2004; Woolner et al., 2008; Wuhr et al., 2008), little evidence demonstrates that Myo X has microtubule based motor activity. Thus, we speculate that microtubule dependent motor protein, kinesin, may be responsible for Myo X anterograde transportation in axons. To this end, KIF13B was identified as a Myo X binding partner to be responsible for Myo X anterograde transportation. Interesting, KIF13B, a kinesin family motor protein, plays an essential role in anterograde transport of PI(3,4,5)P3 (Horiguchi et al., 2006), a binding partner and regulator of Myo X (Figure 4 and Figure 5). Moreover, KIF13B exerts similar functions as Myo X in promoting axon initiation and terminal targeting. In aggregates, our results suggest that Myo X appears to be a cargo of KIF13B during its axonal transportation, and at the same time, Myo X’s actin based motor activity is suppressed by KIF13B.

Netrin-1/DCC signaling is involved in many aspects of axon development, including axon outgrowth and guidance, growth cone steering and axon branching. The canonical model for Netrin-1’s function in axon guidance is that Netrin-1 acts as a long-range diffusible guidance cue, centered in the midline (e.g., floor plate in the developing spinal cord), attracting or repulsing axons for their midline crossing. Recent studies have shown that Netrin-1 is produced not only in the midline, but also in neural progenitor cells (NPCs) in the ventricular zone (VZ), and deposited on the pial surface as a haptotactic adhesive substrate, where it guides DCC^+^ axon growth (Dominici et al., 2017). In developing cerebral cortex, Netrin-1 mRNA is highly expressed in VZ/SVZ (Zhang et al., 2018). These results implicate that Netrin-1/DCC signaling in local microenvironment surrounding new-born neurons is important for axon development. In line with this view, we found that the axon initiation was slowed down by Netrin-1 KO in the local region (Figure 2A and 2B), and Netrin-1 overexpression diminished Myo X-deficiency-induced axon initiation deficit.

In light of our results, we speculate the existence of DCC-Myo X-KIF13B complex. Netrin-1 may increase the complex formation by generating more PI(3,4,5)P3, which binds to Myo X, changes Myo X conformation for DCC and KIF13B binding and then undergoes anterograde transport along microtubules (Figure 7). Myo X’s motor activity may be suppressed by disconnecting Myo X with F-actin filaments, as we can see that KIF13B suppress Myo X motility in dendrite-like actin filaments. In this complex, Myo X acts as a central adapter protein to link its cargos of DCC and PI(3,4,5)P3/PI(3,4)P2 with KIF13B. Such Myo X containing complex may be crucial for Netrin-1 induced axonal outgrowth and growth cone attractive response.

**Figure 7.**
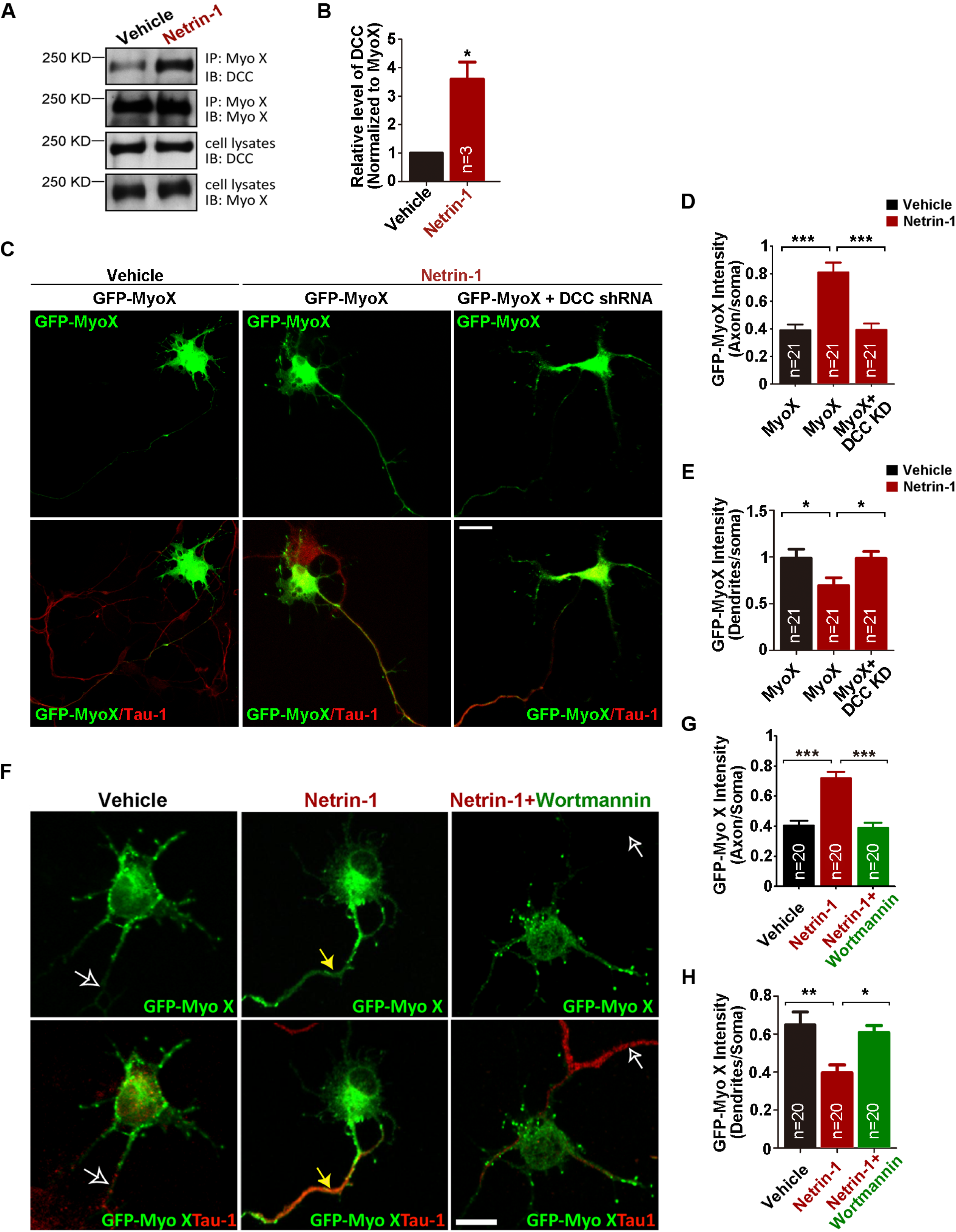
Graphical abstract.

## MATERIALS and METHODS

### Animals

Myo X^f/f^ mice were generated as previously described (Wang et al., 2018) and NTN1^f/f^ mice were generated as illustrated in Figure 2-Figure Supplement 1A. All the mouse lines indicated above were maintained in C57BL/6 background for >6 generations. Timed pregnant female mice were obtained by crossing with male mice overnight, and the noon of the following day was designated as E0.5 if the vaginal plug was detected.

### Reagents

For immunostaining analysis, the following primary antibodies were used: mouse monoclonal anti-Tau-1 (05-838, 1:1000), mouse monoclonal anti-MAP2 (05-346, 1:500), rat monoclonal anti-tubulin (MAB1864, 1:500) from Millipore; chicken polyclonal anti-GFP (ab13970, 1:1000), mouse monoclonal anti-c-Myc (ab32, 1:200) from Abcam. For immunoblotting analysis, the following primary antibodies were used: goat polyclonal anti-DCC (sc-6535, 1:200) from Santa Cruz Biotechnology; rabbit polyclonal anti-KIF13B (PA540807,1:500) from Invitrogen; mouse monoclonal anti-α-tubulin (T5168, 1:4000) and mouse monoclonal anti-GFP (11814460001,1:2000) from Sigma-Aldrich; rabbit polyclonal anti-Myc (ab9106, 1:1000) from Abcam; rabbit polyclonal anti-MyoX was prepared as previously described (Zhu et al., 2017); the polyclonal anti-KIF13B antibody was kindly provided by Dr. Hiroaki Miki (Osaka University, Japan). For immunoprecipitation assay, rabbit polyclonal anti-GFP (A11122) was purchased from Invitrogen. Alexa Fluor 488-, 555- and 647-coupled secondary antibodies against mouse, rat, chicken or goat and HRP-conjugated secondary antibodies against mouse or rabbit were purchased from Jackson ImmunoReseach. Alexa Fluor 350-phalloidin (A22281) and lipofectamine 2000 (11668-019) were obtained from Invitrogen. Wortmannin was from Millipore.

### Expression vectors

The cDNA of mouse *Myo X* was subcloned into mammalian expression vector (pEGFP-C1) fused with GFP at the amino-terminus, as described previously (Zhu et al., 2007). GFP-MyoXΔPH and GFP-MyoXΔFERM were generated with Q5 Site-Directed Mutagenesis Kit (New England Biolabs, E0554S), where the amino acids (1212-1253) and (1799-2058) were deleted, respectively. Myo X-Head, Myo XΔMotor (headless Myo X, hMyo X), Myo X MyTH4-FERM and MyoX FERM were amplified and subcloned into pEGFP-C1. GFP-hMyoXΔPH2 and GFP-hMyoXΔPH3 were generated with Q5 Site-Directed Mutagenesis Kit, where the amino acids (1215-1316) and (1381-1506) were deleted, respectively. Also, GFP-hMyoX (KK1225/6AA) were generated with Q5 Site-Directed Mutagenesis Kit. The cDNA of human KIF13B was subcloned into mammalian expression vector (pRK5) fused with a Myc tag at the amino terminus. KIF13B^1-557^, KIF13B^1-1531^, KIF13B^558-1826^, KIF13B^990-1826^ and KIF13B^1532-1826^ were amplification by PCR and subcloned into pRK5 though corresponding restriction enzymes or exonuclease III. The expression vector s of all GST fusion proteins was constructed by ligation into pGEX-6p-1. mCherry-Myo X was kindly provided by Dr. Staffan Strömblad (Karolinska Institutet, Huddinge, Sweden).

The miRNA expression vectors for Myo X and DCC were generated by the BLOCK-iT Lentiviral miR RNA Expression System (Invitrogen, Carlsbad, CA) according to the manufacturer’s instruction as previously described (Liu et al., 2012; Zhu et al., 2007). The shRNA expression vectors for mouse KIF13B were generated using pll3.7 lentiviral vector, and the target sequences for KIF13B shRNAs are below: 5’- GCAGATAACTATGACGAAACC-3’ (KIF13B shRNA-1); 5’- GGATTTAGCTGGCAGTGAACG-3’ (KIF13B shRNA-2). Netrin-1 shRNA was constructed into pll3.7 lentiviral vector as previously reported and the target sequence was 5’-GGGTGCCCTTCCAGTTCTA-3’ (Chen et al., 2017). In addition, we also generated the RFP-Scramble shRNA and RFP-KIF13B shRNA expression vectors by replacing GFP with RFP in the pll3.7 plasmids. The authenticity of all constructs was verified by DNA sequence.

### Cell cultures and transfections

Primary cortical neurons were cultured as described previously (Zhu et al., 2007). In brief, embryos (E17) were removed from anesthetized pregnant mice. Cerebral cortices were separated and chopped into small pieces. After incubation in 0.125% Trypsin plus with 0.05% DNase in HBSS at 37°C for 20 mins, cells were triturated with fire-polished glass Pasteur pipet and filted with 40 μm filter. Dissociated cells were suspended in DMEM with 10% FBS and plated on poly-D-lysine coated dishes or glass coverslips at 37 °C in a 5% CO2 atmosphere. 4 hours later, the medium was changed into Neurobasal medium with 2% B27 supplement and 2 mM Glutamax. For transfection, neurons were electroporated immediately after dissociation using the Mouse Neuron Nucleofector Kit (Amaxa). In brief, 3×10^6^ neurons were resuspended in 100 µl of nucleofectamine solution containing 3 µg of plasmid and electroporated with Program O-003 of Nucleofector™ II.

NLT cells and HEK293 cells were grown in DMEM supplemented with 10% FBS and 100 units ml^−1^ penicillin-streptomycin. For imaging experiments, 50%-70% confluent NLT cells in 12-well plate were transfected with 1.6 μg indicated plasmids using 3ul lipofectamine in DMEM without FBS and antibiotics. For Western blot and co-immunoprecipitation, HEK293 cells were transfected using polyetherimide (PEI). Stable HEK 293 cell line expressing human netrin-1 was used as described previously (Ren et al., 2004; Xie et al., 2005; Zhu et al., 2007).

### In Utero electroporation

The *in utero* electroporation was carried out as described previously with some modifications (Yu et al., 2012). Briefly, plasmids were microinjected into the lateral cerebral ventricle of E 14.5 or E15.5 mouse embryos through the uterine wall. Then, 35 V square-wave pulse was delivered across the head for 5 times through ECM-830 (BTX, Holliston, MA). Embryos were then allowed to develop to E18.5, P7 or P14. The transfected brains were then fixed with 4% PFA/PBS overnight at 4°C. The brains were sectioned with a freezing microtome at about 50 µm.

### Immunostaining analysis

Cells were fixed in 4% PFA for 10 min at room temperature, permeabilized with 0.1% Triton X-100 for 8 min and blocked in 2% bovine serum albumin for 1 h in 0.01 M phosphate-buffered saline (PBS; pH 7.4). Subsequently, cells were incubated with primary antibodies diluted in the blocking solution for 2 hours and washed three times with PBS. And they were incubated with appropriate fluorochrome-conjugated secondary antibodies for 1 h and washed 3 times.

### Live cell time-lapse imaging and kymography analyses

Transfected neurons were grown on Lab-Tek II Chambered Coverglass (Thermo Fisher Scientific, USA) in DMEM supplemented with 10% FBS and antibiotics. For visualizing GFP-Myo X movement, the Lab-Tek II Chambered Coverglass were then fitted into a temperature-controlled chamber on the microscope stage of LSM 710 confocal laser scanning microscopy (Carl Zeiss, Germany) for observation at 37 °C in a 5% CO2 atmosphere. Time-lapse intervals were 5s and neurons were imaged over periods of 10 minutes. Imagines were acquired with an X63/1.4 N.A. objective at a resolution of 1,024 X 1,024 pixels. The software ImageJ (FIJI) was used for Tracking analysis and Kymographic analysis. In brief, the travelling path and velocity of GFP-Myo X puncta were recorded with the “Manual Tracking” plugin by clicking on the GFP-Myo X puncta on the temporal stacks. For kymographic analysis, a segmented line was used to draw a region of interest (ROI) and then the “KymographBuilder” plugin was used to produce kymographs for the selected segments.

### Fluorescence recovery after photobleaching

The experiments were performed using the LSM 710 confocal laser scanning microscopy. Imagines were acquired with an X63/1.4 N.A. objective at a resolution of 1,024 X 1,024 pixels. A region at the proximal axon was bleached with high laser power and fluorescence recovery was observed for a period of 10min. For FRAP analysis, the mean intensity of the bleached area was normalized with the initial fluorescence intensity before bleaching.

### Protein–protein interaction assays

Immunoprecipitation was carried out as previously described (Ren et al., 2001). Cell lysates (1 mg protein) were incubated at 4 °C for 6 hours with the indicated antibodies (1–2 μg) in a final volume of 1 mL modified RIPA lysis buffer with protease inhibitors. After the addition of protein A-G-agarose beads, each reaction was incubated at 4 °C for 1 h. The immunoprecipitate complexes were collected by centrifugation and washed 3 times with washing buffer (20 mM Tris-HCl, 10 mM NaCl, 1 mM EDTA, 0.5% NP40). Immune complexes were resolved by SDS–PAGE and subjected to immunoblotting. GST pulldown assay was carried out as described previously (Ren et al., 2001). Transiently transfected HEK 293 cells were lysed in the modified RIPA buffer (50 mM Tris-HCl, pH 7.4, 150 mM sodium chloride, 1% NP-40, 0.25% sodium-deoxycholate, and proteinase inhibitors). Cell lysates were precleared with GST immobilized on glutathione–Sepharose 4B (GE Healthcare) and then incubated with the indicated GST fusion proteins (2–5 μg) immobilized on glutathione–Sepharose beads at 4°C overnight with constant rocking. The beads were washed three times with modified RIPA buffer, and bound proteins were resolved by SDS–PAGE and subjected to immunoblotting.

### Imaging quantification and statistical analyses

Immunostaining sections and cells were observed under a Zeiss LSM 710 confocal microscope with ZEN 2012 software, and only the brightness, contrast, and color balance were optimized after imaging. The software ImageJ was used to measure fluorescence intensity in all fixed images. The software ImageJ was used to measure fluorescence intensity in all fixed images. In brief, the RGB images were converted into 8-bit grayscale images and inverted to negative images for analysis. After converted to uncalibrated optical density value, the area of axons, soma and dendrites was selected with ROI tools and calculated. GFP-MyoX intensity in axons or dendrites was normalized by that in its soma regions. Statistical analyses were performed using either unpaired 2-tailed Student’s t test or 1-way analysis of variance (ANOVA) followed by a protected least significant difference Fisher’s post hoc test for multiple comparisons. Statistical evaluations were performed with the software Graph Pad Prism version 5.0. The data are presented as the mean ± standard error of the mean (SEM). P values less than 0.05 were considered significant.

## Acknowledgement

We are grateful to Dr. Hiroaki Miki (Osaka University, Japan) for providing KIF13B reagents. This project is supported by National Institute of Aging, NIH (AG045781).

---

**Figure 1-Figure Supplement 1.**
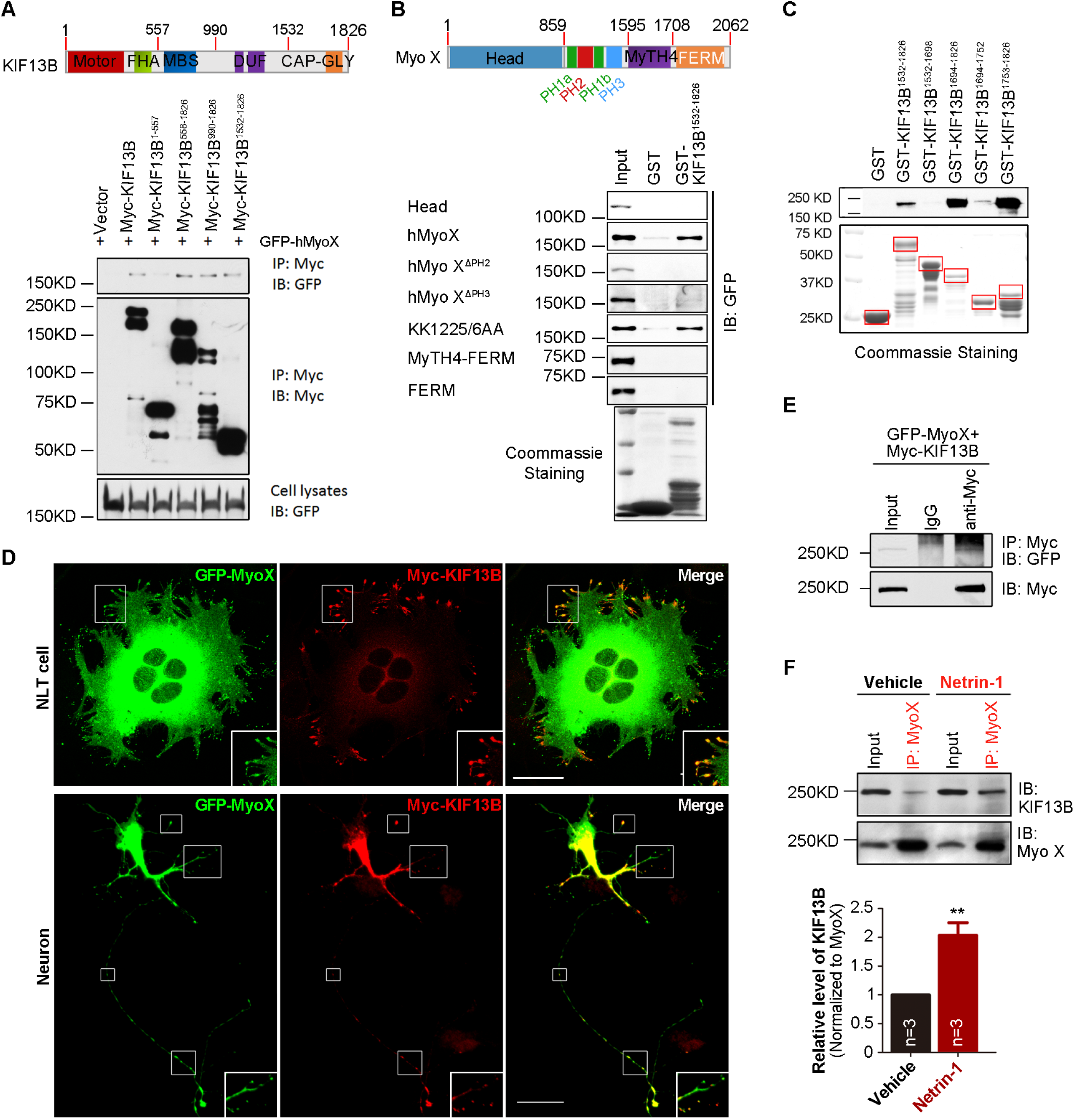
Defective axon ipsilateral branching in Myo X-KO cortical neurons. **(A)** Representative images of P7 cerebral cortex electroporated with GFP (Control) or Cre-GFP (Myo X KO) plasmids at E15.5. Scale Bar=500μm. **(B)** Quantification of axon ipsilateral branching. Student’s t test, p =0.0356. **(C)** Representative images of P3 cerebral cortex electroporated with MyoX miRNA (Myo X KD) or control miRNA (Control) at E15.5. Scale Bar=200μm. **(D)** Axon distribution was illustrated and quantified. Student’s t test, p<0.0001. Data are presented as the means ± SEM. The numbers of brain sections scored are from 3 different brains for each group and indicated on the graphs. **, P<0.01; ***, P<0.001.

**Figure 2-Figure Supplement 1.**
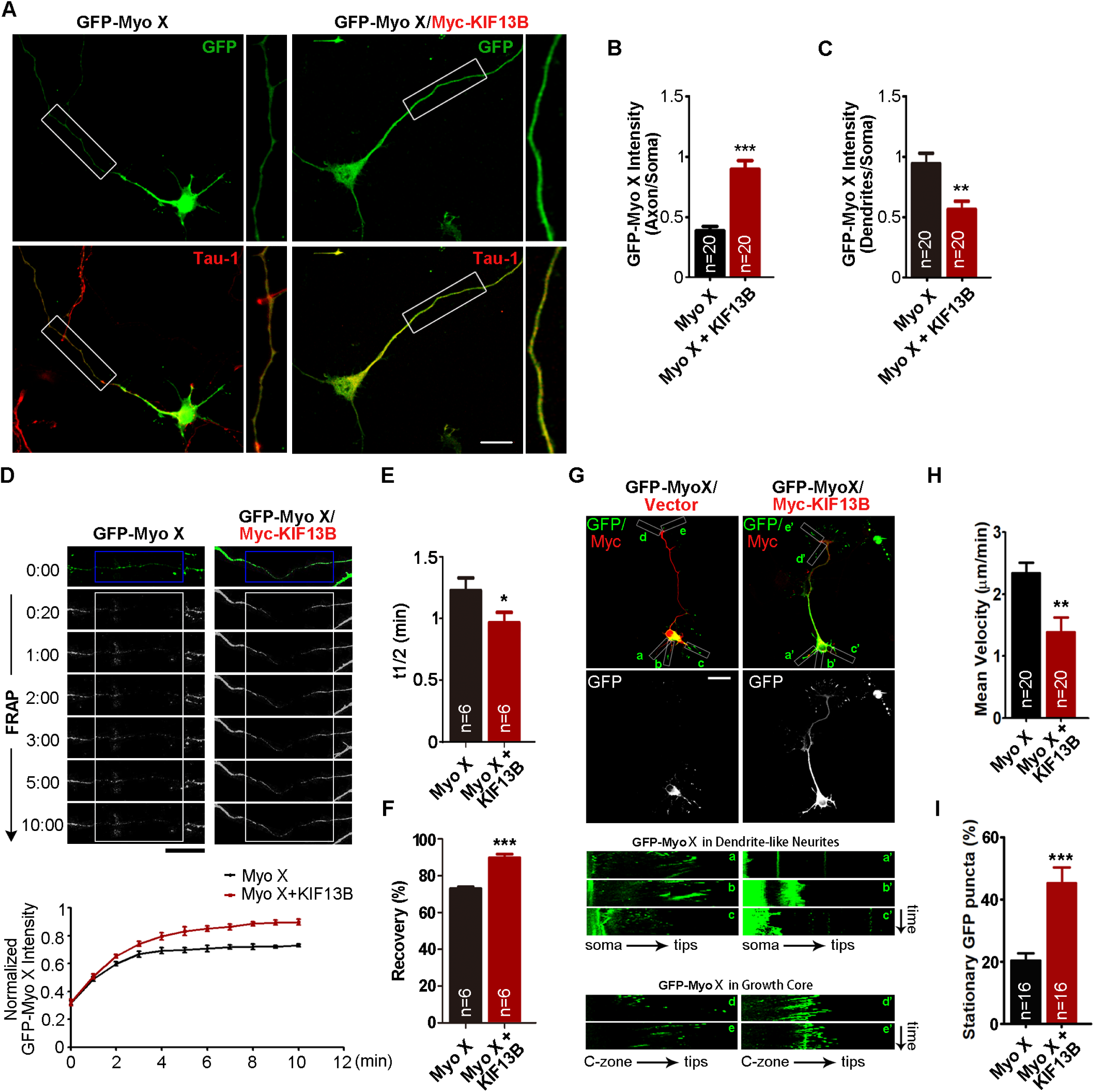
Axon ipsilateral branching in Netrin-1 and DCC-KO cortical neurons. **(A)** Diagram of how to generate NTN1^f/f^ and NTN1 CKO mice. **(B-C)** Representative images of P14 cerebral cortex electroporated with Netrin-1 shRNA, DCC miRNA or Control plasmids at E15.5 as well as quantification of axon ipsilateral branching. Scale Bar=100μm. Student’s t test, for Netrin-1 KD group, p=0.9206, for DCC KD group, p=0.0155. Data are presented as the means ± SEM. The numbers of brain sections scored are from 3 different brains for each group and indicated on the graphs. ns, no significant difference; *, P<0.05.

**Figure 3-Figure Supplement 1.**
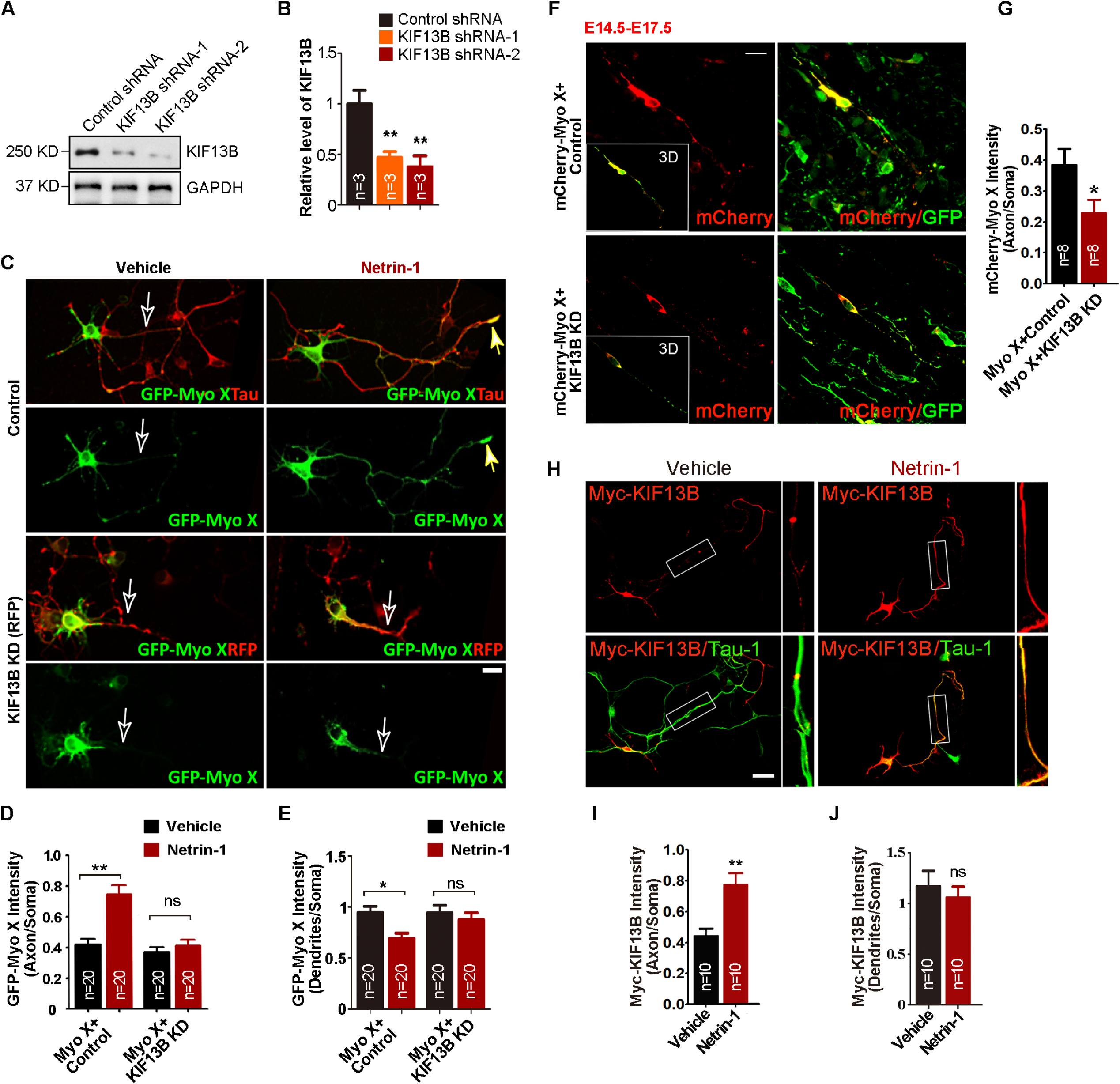
Requirement of Myo X’s PH and FERM domains for Netrin-1 increase of axonal distribution of exogenous GFP-Myo X. **(A)** Neurons transfected with GFP-Myo X and its deletion mutants (illustrated in the left panels) were treated with vehicle or Netrin-1 and then stained with anti-Tau-1 antibody at DIV 3. Scale Bar=10μm. **(B)** Quantification of axonal distribution of GFP-Myo X or its deletion mutants. Student’s t test, for Myo X group, p =0.0003, for Myo Δ PH group, p=0.8293, for Myo Δ FERM group, p=0.4571. Data are presented as the means ± SEM. The numbers of neurons scored in these groups are from 3 different experiments and indicated on the graphs. ns, no significant difference; ***, P<0.001.

**Figure 3-Figure Supplement 2.**
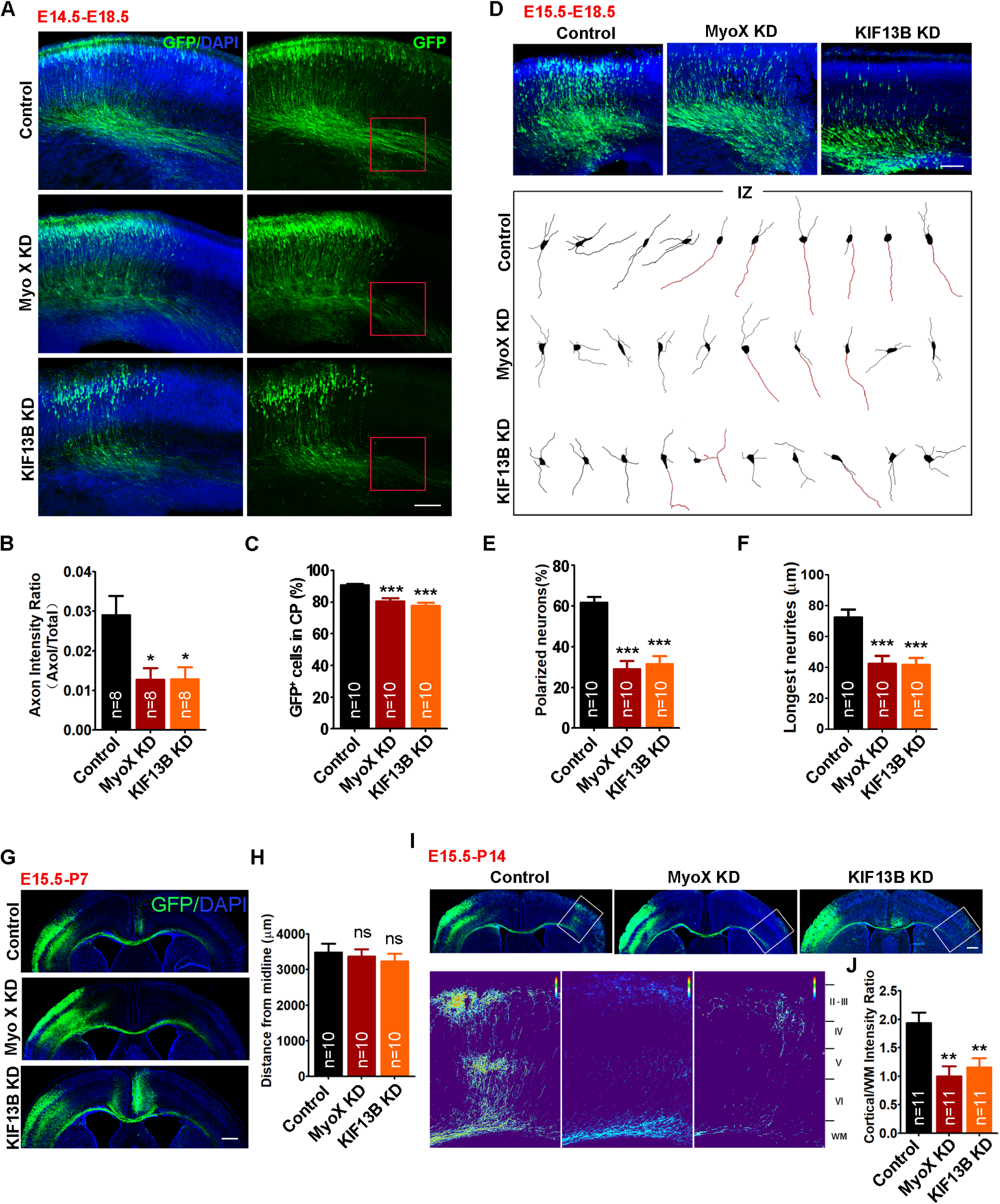
DCC and PI3K activity dependent axonal distribution of GFP-Myo X in response to Netrin-1. **(A)** Immunoprecipitation analysis of DCC association with Myo X in cultured cortical neurons. **(B)** Quantifications of DCC binding with MyoX. student’s t test, p = 0.0122. **(C)** Neurons transfected with GFP-Myo X or GFP-Myo X together with DCC miRNA were treated with vehicle or Netrin-1 and then stained with anti-Tau-1 antibody at DIV 3. Scale Bar=10μm. **(D)** Quantification of axonal distribution of GFP-Myo X. One-way ANOVA, for Myo X group with vehicle stimulation and Myo X group with Netrin-1 stimulation, p <0.0001; for Myo X and Myo X+DCC KD groups, p <0.0001. **(E)** Quantification of dendritic distribution of GFP-Myo X. One-way ANOVA, for Myo X groups with vehicle and Myo X group with Netrin-1 stimulation, p=0.0305, for Myo X and Myo X+DCC KD groups, p=0.0318. **(F)** GFP-Myo X-expressing neurons were treated with vehicle and Netrin-1 with or without PI3 kinase inhibitor, wortmannin (10 nM) for 1 hr and subjected to immunostaining with anti-Tau-1 antibody at DIV 3. Scale Bar=10μm. **(G)** Quantification of axonal distribution of GFP-Myo X. One-way ANOVA, p<0.0001 for both groups. **(H)** Quantification of dendritic distribution of GFP-Myo X. One-way ANOVA, for vehicle and Nerin-1 groups, p=0.0023; for Netrin-1 and Netrin-1+wortmannin groups, p=0.0124. Data are presented as the means ± SEM. The numbers of brain sections scored are from 3 different brains for each group and indicated on the graphs. ns, no significant difference; *, P<0.05; **, P<0.01; ***, P<0.001.

**Figure 5-Figure Supplement 1.**
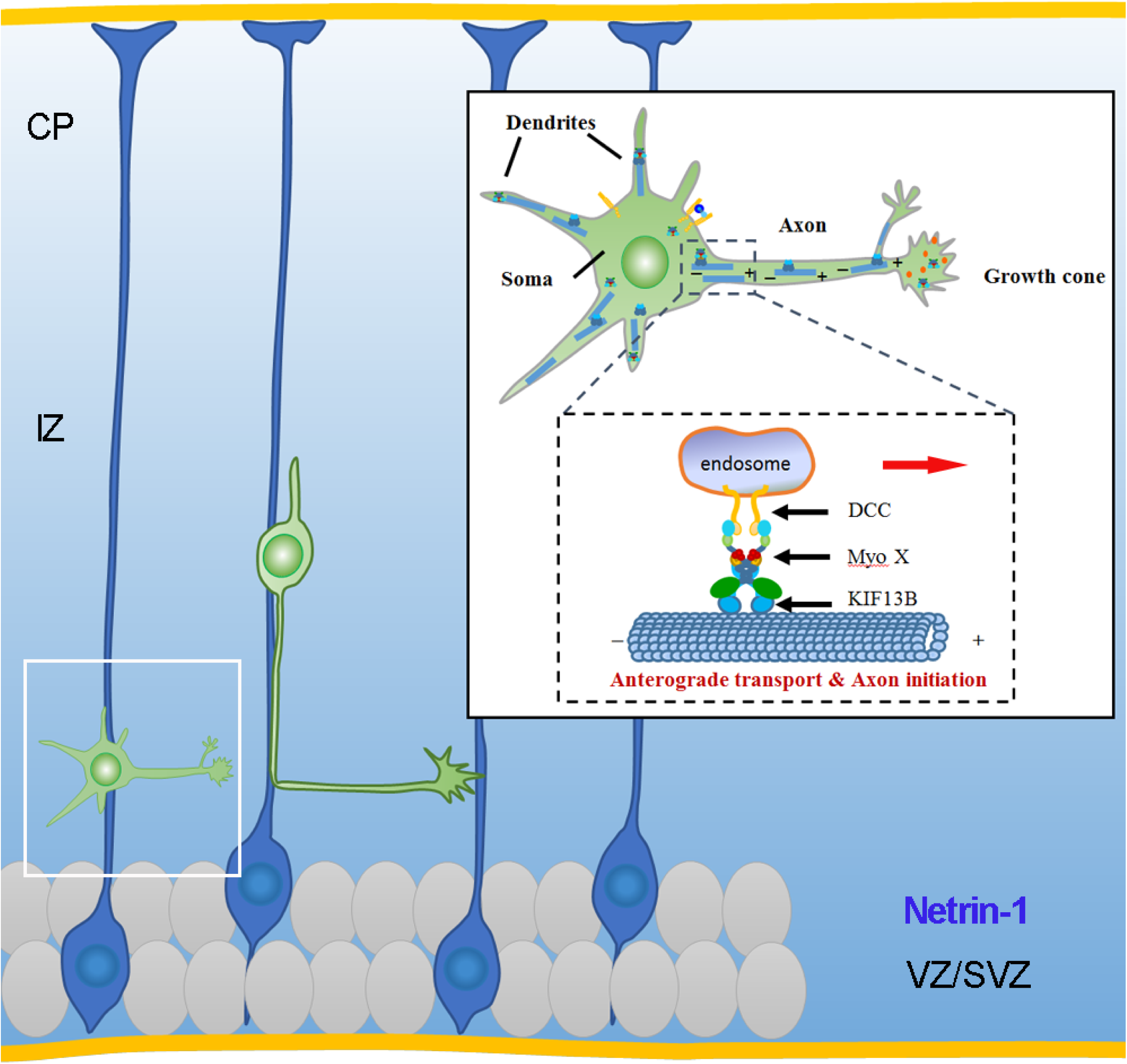
Requirement of KIF13B for Netrin-1 induced axonal distribution of GFP-Myo X. **(A)** Western blot showing the silence effect of KIF13B shRNA in cultured cortical neurons. **(B)** Quantification of KIF13B knockdown efficiency. One-way ANOVA, p=0.0036 for KIF13B shRNA-1 group; p=0.0015 for KIF13B shRNA-2 group. **(C)** Neurons transfected with GFP-Myo X together with control shRNA or KIF13B shRNA respectively were treated with vehicle or Netrin-1 for 1 hr and subjected to immunostaining at DIV 3. Scale Bar=10μm. **(D)** Quantification of GFP-Myo X intensity in axons. The axonal GFP-Myo X level was normalized to somatic GFP-Myo X. One-way ANOVA, p<0.0001 between GFP-Myo X+Control groups; p=0.9059 between GFP-Myo X+KIF13B KD groups. **(E)** Quantification of GFP-Myo X intensity in dendrites. The dendritic GFP-Myo X level was normalized to somatic GFP-Myo X. One-way ANOVA, p=0.0293 between GFP-Myo X+Control groups; p=0.8791 between GFP-Myo X+KIF13B KD groups. **(F)** Representative images of cortical neurons electroporated with indicated plasmids at E15.5. Scale Bar=10μm. **(G)** Quantification of mCherry-Myo X intensity in axons. The axonal mCherry-Myo X level was normalized to somatic mCherry-Myo X. Student’s t test, p=0.0331. **(H)** Neurons transfected with Myc-KIF13B were treated with vehicle or Netrin-1 for 1 hr and subjected to immunostaining with indicated antibodies at DIV 3. Images marked in rectangulars were amplified and showed in the right panels. Scale Bar=20μm. **(I)** Quantification of Myc-KIF13B intensity in axons. The axonal Myc-KIF13B level was normalized to somatic Myc-KIF13B. Student’s t test, p=0.0026. **(J)** Quantification of Myc-KIF13B intensity in dendrites. The dendritic Myc-KIF13B level was normalized to somatic Myc-KIF13B. Student’s t test, p=0.591. Data are presented as the means ± SEM. The numbers of neurons scored in these groups are from 3 different experiments and indicated on the graphs. ns, no significant difference; *, P<0.05; **, P<0.01; ***, P<0.001.

## Reference

1. Ackerman, S.L., Kozak, L.P., Przyborski, S.A., Rund, L.A., Boyer, B.B., and Knowles, B.B. (1997). The mouse rostral cerebellar malformation gene encodes an UNC-5-like protein. Nature 386, 838–842.

2. Alcamo, E.A., Chirivella, L., Dautzenberg, M., Dobreva, G., Farinas, I., Grosschedl, R., and McConnell, S.K. (2008). Satb2 regulates callosal projection neuron identity in the developing cerebral cortex. Neuron 57, 364–377.

3. Barnes, A.P., and Polleux, F. (2009). Establishment of axon-dendrite polarity in developing neurons. Annual review of neuroscience 32, 347–381.

4. Berg, J.S., and Cheney, R.E. (2002). Myosin-X is an unconventional myosin that undergoes intrafilopodial motility. Nature cell biology 4, 246–250.

5. Berg, J.S., Derfler, B.H., Pennisi, C.M., Corey, D.P., and Cheney, R.E. (2000). Myosin-X, a novel myosin with pleckstrin homology domains, associates with regions of dynamic actin. Journal of cell science 113 Pt 19, 3439–3451.

6. Braisted, J.E., Catalano, S.M., Stimac, R., Kennedy, T.E., Tessier-Lavigne, M., Shatz, C.J., and O’Leary, D.D. (2000). Netrin-1 promotes thalamic axon growth and is required for proper development of the thalamocortical projection. The Journal of neuroscience 20, 5792–5801.

7. Brown, A. (2003). Axonal transport of membranous and nonmembranous cargoes: a unified perspective. The Journal of cell biology 160, 817–821.

8. Campbell, D.S., and Holt, C.E. (2003). Apoptotic pathway and MAPKs differentially regulate chemotropic responses of retinal growth cones. Neuron 37, 939–952.

9. Chen, J.Y., He, X.X., Ma, C., Wu, X.M., Wan, X.L., Xing, Z.K., Pei, Q.Q., Dong, X.P., Liu, D.X., Xiong, W.C., et al. (2017). Netrin-1 promotes glioma growth by activating NF-kappaB via UNC5A. Scientific reports 7, 5454.

10. Colamarino, S.A., and Tessier-Lavigne, M. (1995). The role of the floor plate in axon guidance. Annual review of neuroscience 18, 497–529.

11. Cox, D., Berg, J.S., Cammer, M., Chinegwundoh, J.O., Dale, B.M., Cheney, R.E., and Greenberg, S. (2002). Myosin X is a downstream effector of PI(3)K during phagocytosis. Nature cell biology 4, 469–477.

12. Culotti, J.G., and Merz, D.C. (1998). DCC and netrins. Current opinion in cell biology 10, 609–613.

13. Dent, E.W., and Gertler, F.B. (2003). Cytoskeletal dynamics and transport in growth cone motility and axon guidance. Neuron 40, 209–227.

14. Dominici, C., Moreno-Bravo, J.A., Puiggros, S.R., Rappeneau, Q., Rama, N., Vieugue, P., Bernet, A., Mehlen, P., and Chedotal, A. (2017). Floor-plate-derived netrin-1 is dispensable for commissural axon guidance. Nature 545, 350–354.

15. Forcet, C., Stein, E., Pays, L., Corset, V., Llambi, F., Tessier-Lavigne, M., and Mehlen, P. (2002). Netrin-1-mediated axon outgrowth requires deleted in colorectal cancer-dependent MAPK activation. Nature 417, 443–447.

16. Hedgecock, E.M., Culotti, J.G., and Hall, D.H. (1990). The unc-5, unc-6, and unc-40 genes guide circumferential migrations of pioneer axons and mesodermal cells on the epidermis in C. elegans. Neuron 4, 61–85.

17. Hirokawa, N., Niwa, S., and Tanaka, Y. (2010). Molecular motors in neurons: transport mechanisms and roles in brain function, development, and disease. Neuron 68, 610–638.

18. Hong, K., Hinck, L., Nishiyama, M., Poo, M.M., Tessier-Lavigne, M., and Stein, E. (1999). A ligand-gated association between cytoplasmic domains of UNC5 and DCC family receptors converts netrin-induced growth cone attraction to repulsion. Cell 97, 927–941.

19. Horiguchi, K., Hanada, T., Fukui, Y., and Chishti, A.H. (2006). Transport of PIP3 by GAKIN, a kinesin-3 family protein, regulates neuronal cell polarity. The Journal of cell biology 174, 425–436.

20. Hwang, Y.S., Luo, T., Xu, Y., and Sargent, T.D. (2009). Myosin-X is required for cranial neural crest cell migration in Xenopus laevis. Developmental dynamics 238, 2522–2529.

21. Keleman, K., and Dickson, B.J. (2001). Short- and long-range repulsion by the Drosophila Unc5 netrin receptor. Neuron 32, 605–617.

22. Kerber, M.L., and Cheney, R.E. (2011). Myosin-X: a MyTH-FERM myosin at the tips of filopodia. Journal of cell science 124, 3733–3741.

23. Lai, M., Guo, Y., Ma, J., Yu, H., Zhao, D., Fan, W., Ju, X., Sheikh, M.A., Malik, Y.S., Xiong, W., et al. (2015). Myosin X regulates neuronal radial migration through interacting with N-cadherin. Frontiers in cellular neuroscience 9, 326.

24. Leonardo, E.D., Hinck, L., Masu, M., Keino-Masu, K., Ackerman, S.L., and Tessier-Lavigne, M. (1997). Vertebrate homologues of C. elegans UNC-5 are candidate netrin receptors. Nature 386, 833–838.

25. Li, W., Lee, J., Vikis, H.G., Lee, S.H., Liu, G., Aurandt, J., Shen, T.L., Fearon, E.R., Guan, J.L., Han, M., et al. (2004). Activation of FAK and Src are receptor-proximal events required for netrin signaling. Nature neuroscience 7, 1213–1221.

26. Li, X., Saint-Cyr-Proulx, E., Aktories, K., and Lamarche-Vane, N. (2002). Rac1 and Cdc42 but not RhoA or Rho kinase activities are required for neurite outgrowth induced by the Netrin-1 receptor DCC (deleted in colorectal cancer) in N1E-115 neuroblastoma cells. The Journal of biological chemistry 277, 15207–15214.

27. Liu, G., Beggs, H., Jurgensen, C., Park, H.T., Tang, H., Gorski, J., Jones, K.R., Reichardt, L.F., Wu, J., and Rao, Y. (2004). Netrin requires focal adhesion kinase and Src family kinases for axon outgrowth and attraction. Nature neuroscience 7, 1222–1232.

28. Liu, Y., Peng, Y., Dai, P.G., Du, Q.S., Mei, L., and Xiong, W.C. (2012). Differential regulation of myosin X movements by its cargos, DCC and neogenin. Journal of cell science 125, 751–762.

29. Maday, S., Twelvetrees, A.E., Moughamian, A.J., and Holzbaur, E.L. (2014). Axonal transport: cargo-specific mechanisms of motility and regulation. Neuron 84, 292–309.

30. Ming, G., Song, H., Berninger, B., Inagaki, N., Tessier-Lavigne, M., and Poo, M. (1999). Phospholipase C-gamma and phosphoinositide 3-kinase mediate cytoplasmic signaling in nerve growth cone guidance. Neuron 23, 139–148.

31. Ming, G.L., Wong, S.T., Henley, J., Yuan, X.B., Song, H.J., Spitzer, N.C., and Poo, M.M. (2002). Adaptation in the chemotactic guidance of nerve growth cones. Nature 417, 411–418.

32. Namba, T., Kibe, Y., Funahashi, Y., Nakamuta, S., Takano, T., Ueno, T., Shimada, A., Kozawa, S., Okamoto, M., Shimoda, Y., et al. (2014). Pioneering axons regulate neuronal polarization in the developing cerebral cortex. Neuron 81, 814–829.

33. Nie, S., Kee, Y., and Bronner-Fraser, M. (2009). Myosin-X is critical for migratory ability of Xenopus cranial neural crest cells. Developmental biology 335, 132–142.

34. Pi, X., Ren, R., Kelley, R., Zhang, C., Moser, M., Bohil, A.B., Divito, M., Cheney, R.E., and Patterson, C. (2007). Sequential roles for myosin-X in BMP6-dependent filopodial extension, migration, and activation of BMP receptors. The Journal of cell biology 179, 1569–1582.

35. Plantard, L., Arjonen, A., Lock, J.G., Nurani, G., Ivaska, J., and Stromblad, S. (2010). PtdIns(3,4,5)P(3) is a regulator of myosin-X localization and filopodia formation. Journal of cell science 123, 3525–3534.

36. Ren, X.R., Du, Q.S., Huang, Y.Z., Ao, S.Z., Mei, L., and Xiong, W.C. (2001). Regulation of CDC42 GTPase by proline-rich tyrosine kinase 2 interacting with PSGAP, a novel pleckstrin homology and Src homology 3 domain containing rhoGAP protein. The Journal of cell biology 152, 971–984.

37. Ren, X.R., Ming, G.L., Xie, Y., Hong, Y., Sun, D.M., Zhao, Z.Q., Feng, Z., Wang, Q., Shim, S., Chen, Z.F., et al. (2004). Focal adhesion kinase in netrin-1 signaling. Nature neuroscience 7, 1204–1212.

38. Shekarabi, M., and Kennedy, T.E. (2002). The netrin-1 receptor DCC promotes filopodia formation and cell spreading by activating Cdc42 and Rac1. Molecular and cellular neurosciences 19, 1–17.

39. Tokuo, H., and Ikebe, M. (2004). Myosin X transports Mena/VASP to the tip of filopodia. Biochemical and biophysical research communications 319, 214–220.

40. Tokuo, H., Mabuchi, K., and Ikebe, M. (2007). The motor activity of myosin-X promotes actin fiber convergence at the cell periphery to initiate filopodia formation. The Journal of cell biology 179, 229–238.

41. Umeki, N., Jung, H.S., Sakai, T., Sato, O., Ikebe, R., and Ikebe, M. (2011). Phospholipid-dependent regulation of the motor activity of myosin X. Nature structural & molecular biology 18, 783–788.

42. Wang, B., Pan, J.X., Yu, H., Xiong, L., Zhao, K., Xiong, S., Guo, J.P., Lin, S., Sun, D., Zhao, L., Guo, H., Mei, L., and Xiong, W.C. (2019). Lack of Myosin X Enhances Osteoclastogenesis and Increases Cell Surface Unc5b in Osteoclast-Lineage Cells. Journal of bone and mineral research 34:939–954.

43. Weber, K.L., Sokac, A.M., Berg, J.S., Cheney, R.E., and Bement, W.M. (2004). A microtubule-binding myosin required for nuclear anchoring and spindle assembly. Nature 431, 325–329.

44. Winkle, C.C., McClain, L.M., Valtschanoff, J.G., Park, C.S., Maglione, C., and Gupton, S.L. (2014). A novel Netrin-1-sensitive mechanism promotes local SNARE-mediated exocytosis during axon branching. The Journal of cell biology 205, 217–232.

45. Woolner, S., O’Brien, L.L., Wiese, C., and Bement, W.M. (2008). Myosin-10 and actin filaments are essential for mitotic spindle function. The Journal of cell biology 182, 77–88.

46. Wuhr, M., Mitchison, T.J., and Field, C.M. (2008). Mitosis: new roles for myosin-X and actin at the spindle. Current biology 18, R912–914.

47. Xie, Y., Ding, Y.Q., Hong, Y., Feng, Z., Navarre, S., Xi, C.X., Zhu, X.J., Wang, C.L., Ackerman, S.L., Kozlowski, D., et al. (2005). Phosphatidylinositol transfer protein-alpha in netrin-1-induced PLC signalling and neurite outgrowth. Nature cell biology 7, 1124–1132.

48. Xie, Y., Hong, Y., Ma, X.Y., Ren, X.R., Ackerman, S., Mei, L., and Xiong, W.C. (2006). DCC-dependent phospholipase C signaling in netrin-1-induced neurite elongation. The Journal of biological chemistry 281, 2605–2611.

49. Yogev, S., and Shen, K. (2017). Establishing Neuronal Polarity with Environmental and Intrinsic Mechanisms. Neuron 96, 638–650.

50. Yonezawa, S., Kimura, A., Koshiba, S., Masaki, S., Ono, T., Hanai, A., Sonta, S., Kageyama, T., Takahashi, T., and Moriyama, A. (2000). Mouse myosin X: molecular architecture and tissue expression as revealed by northern blot and in situ hybridization analyses. Biochemical and biophysical research communications 271, 526–533.

51. Yoshimura, Y., Terabayashi, T., and Miki, H. (2010). Par1b/MARK2 phosphorylates kinesin-like motor protein GAKIN/KIF13B to regulate axon formation. Molecular and cellular biology 30, 2206–2219.

52. Yu, H., Wang, N., Ju, X., Yang, Y., Sun, D., Lai, M., Cui, L., Sheikh, M.A., Zhang, J., Wang, X., et al. (2012). PtdIns (3,4,5) P3 recruitment of Myo10 is essential for axon development. PloS one 7, e36988.

53. Zhang, H., Berg, J.S., Li, Z., Wang, Y., Lang, P., Sousa, A.D., Bhaskar, A., Cheney, R.E., and Stromblad, S. (2004). Myosin-X provides a motor-based link between integrins and the cytoskeleton. Nature cell biology 6, 523–531.

54. Zhang, J.H., Zhao, Y.F., He, X.X., Zhao, Y., He, Z.X., Zhang, L., Huang, Y., Wang, Y.B., Hu, L., Liu, L., et al. (2018). DCC-Mediated Dab1 Phosphorylation Participates in the Multipolar-to-Bipolar Transition of Migrating Neurons. Cell reports 22, 3598–3611.

55. Zhu, X.J., Wang, C.Z., Dai, P.G., Xie, Y., Song, N.N., Liu, Y., Du, Q.S., Mei, L., Ding, Y.Q., and Xiong, W.C. (2007). Myosin X regulates netrin receptors and functions in axonal path-finding. Nature cell biology 9, 184–192.

